# Skp1 proteins are structural components of the synaptonemal complex in *C. elegans*

**DOI:** 10.1101/2023.05.13.540652

**Authors:** Joshua Blundon, Brenda Cesar, Jung Woo Bae, Ivana Čavka, Jocelyn Haversat, Jonas Ries, Simone Köhler, Yumi Kim

**Affiliations:** Department of Biology, Johns Hopkins University, Baltimore, MD 21218, USA; The European Molecular Biology Laboratory, Heidelberg, Germany; Collaboration for joint PhD degree between EMBL and Heidelberg University, Faculty of Biosciences, Heidelberg, Germany

## Abstract

The synaptonemal complex (SC) is a hallmark of meiotic prophase that plays a crucial role in regulating crossovers between homologous chromosomes. Here, we demonstrate that two Skp1-related proteins in *C. elegans*, SKR-1 and SKR-2, serve as structural components of the SC, independent of their canonical functions within the Skp1-Cul1-F-box (SCF) ubiquitin ligase complex. SKR-1 and SKR-2 localize to the central region of the SC, and synapsis requires their dimerization through a hydrophobic interface that overlaps with the binding sites for CUL-1 and F-box proteins. Using *in vitro* reconstitution and *in vivo* analysis of mutant proteins, we show that SKR proteins interact with the other SC proteins using their C-terminal helices to form a soluble complex, which likely represents a basic building block for SC assembly. Our findings demonstrate how conserved Skp1 proteins are repurposed as part of the SC and may provide insight into how synapsis is coupled to cell cycle progression.

## Introduction

Accurate chromosome segregation during meiosis requires that chromosomes pair and undergo crossover recombination with their homologous partners. In most eukaryotes, pairwise interactions between homologs are held together by a zipper-like protein structure known as the synaptonemal complex (SC).^1^ First described by electron microscopy more than 65 years ago, the SC is a defining feature of meiotic chromosomes.^2, 3^ The SC interacts with proteins that promote crossovers and allows them to diffuse and concentrate along the length of the chromosomes, helping to control the number and distribution of crossovers.^4–8^ The SC is conventionally viewed as a tripartite structure consisting of two parallel chromosome axes that organize chromatin into loops and a central region that links the paired homologs.^9^ Although the appearance of the SC is highly similar across eukaryotes, the components comprising the SC have diverged extensively.

Despite the divergence in SC components, its assembly and disassembly are largely governed by post-translational modifications added by conserved cell cycle regulators.^10^ An emerging example is the Skp1-Cul1-F-box (SCF) ubiquitin ligase that regulates synapsis and crossover formation across diverse eukaryotes.^11–16^ The SCF complex is the archetypal cullin-RING ubiquitin ligase,^17^ in which Cul1 serves as the core scaffolding subunit to link a RING domain protein Rbx1 and an adaptor Skp1 (**Figure 1A**).^18^ Rbx1 recruits a ubiquitin-conjugating enzyme (E2) to form the catalytic core, and Skp1 interacts with F-box proteins that recruit a diverse group of substrates to the SCF complex.^19, 20^ By bringing E2 close to substrates, the SCF complex promotes the transfer of ubiquitin to the target protein, and polyubiquitinated substrates are ultimately degraded by the proteasome.^21^

**Figure 1.**
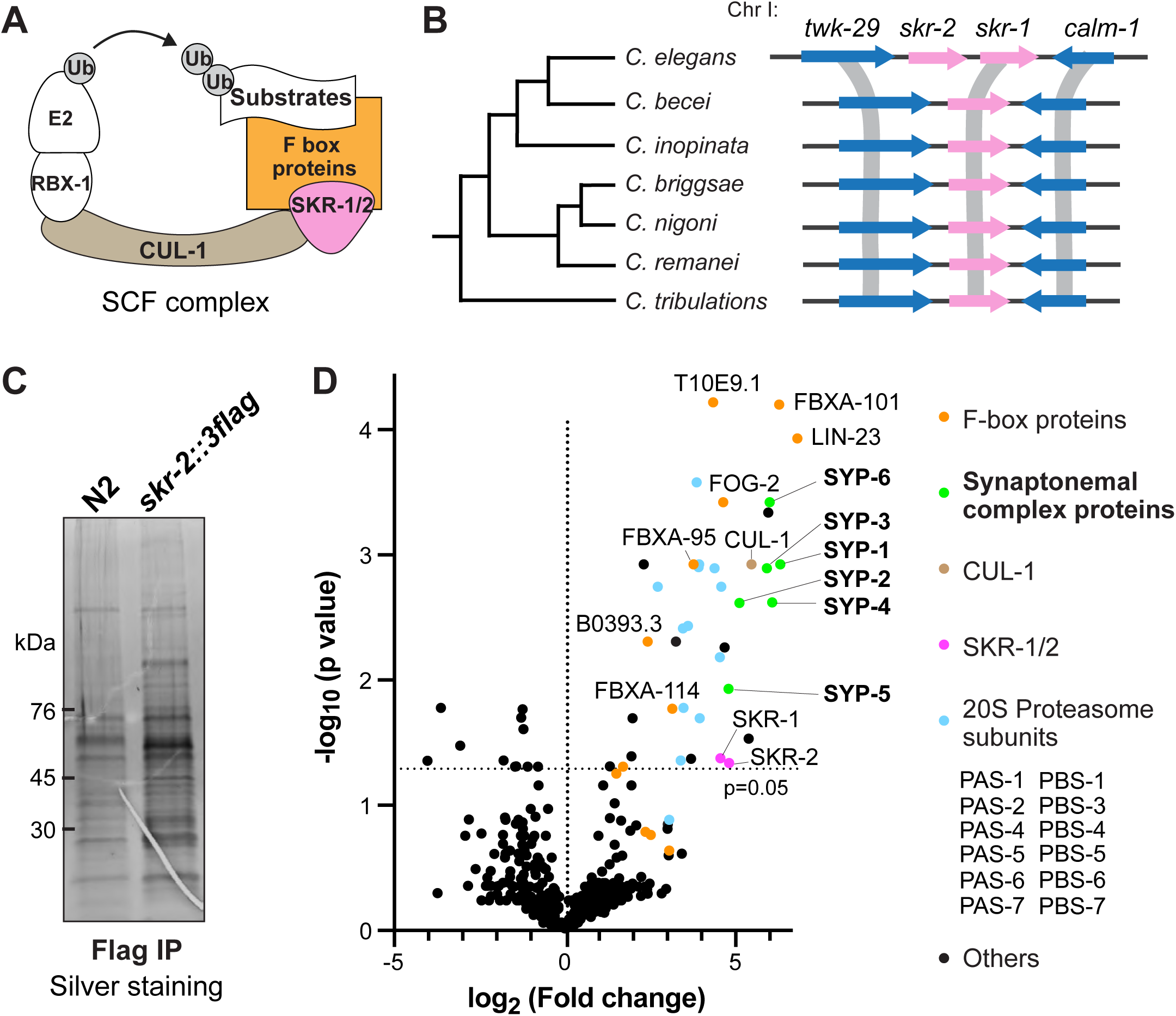
Two paralogous Skp1-related proteins in *C. elegans*, SKR-1 and SKR-2, associate with SC proteins. A) Schematic showing polyubiquitination of a substrate by an E2 ubiquitin-conjugating enzyme and the SCF ubiquitin ligase complex. (B) Synteny analysis of *skr-1* and *skr-2* genes in related *Caenorhabditis* species. Given the requirement of *skr-1* during development,^54^ we infer that *skr-1* has been derived from the ancestral gene. (C) A silver stained SDS-PAGE gel showing purified SKR-2-containing protein complexes using anti Flag beads from a worm strain expressing SKR-2::3Flag. N2 is used as a control. (D) A volcano plot showing proteins enriched in SKR-2 immunoprecipitates over N2 control. Normalized weighted spectra were transformed to logarithmic values (base 2) and analyzed by multiple unpaired t-test. The logarithm of fold change (base 2) is plotted on the x-axis, and the negative logarithm of the p-value (base 10) is plotted on the y-axis. The p-value of 0.05 is indicated by a horizontal dotted line. Proteins enriched in SKR-2 immunoprecipitates are found in the upper right corner of the plot. F-box proteins (orange), SC proteins (green), CUL-1 (brown), SKR-1/2 (magenta), Proteasome subunits (cyan), and other proteins (black) are highlighted and listed on the right.

The function of SCF complexes in meiosis was first discovered through studies in *Arabidopsis,* where ASK1 (*Arabidopsis* Skp1-like) is essential for male meiosis.^22^ ASK1 facilitates nuclear reorganization during early meiotic prophase, which is necessary for homolog pairing and synapsis.^14, 15, 23^ Studies in other species including budding yeast, *Drosophila*, and mice have revealed that depletion of SCF subunits during meiosis leads to severe synapsis defects,^11–13^ indicating the conserved role of SCF in synapsis. In mice, SKP1 localizes to synapsed chromosome axes and controls sister chromatid cohesion, recruitment of meiotic DNA break-forming machinery, and pachytene exit.^11, 24^ Even in fission yeast, which does not form the SC,^25^ Skp1 and an F-box protein are required for processing recombination intermediates.^26^ Several F-box proteins have been demonstrated to play crucial roles in homolog pairing, synapsis/desynapsis, and meiotic recombination across diverse species, and their substrates have started to emerge.^11, 12, 27–32^ However, the list is nowhere near complete, and it is unknown how the SCF complex recognizes a variety of substrates to regulate meiotic chromosome dynamics.

In *C. elegans,* two paralogous Skp1-related (SKR) proteins, SKR-1 and SKR-2, control cell cycle progression during germline development.^16, 33^ During the transition from mitosis to meiosis, SCF complexes containing an F-box protein PROM-1 (SCF^PROM-1^) target a PP2C phosphatase PPM-1.D for degradation, leading to activation of CHK-2 kinase and initiation of meiotic chromosome dynamics.^34–36^ The SCF complex also controls pachytene exit in *C. elegans* and mediates the degradation of translational regulators for maturing oocytes.^37, 38^ Interestingly, SKR-1/2-depleted germlines exhibit an extended region of leptotene/zygotene stage nuclei,^16^ which is now known to reflect a delayed meiotic progression in response to synapsis defects.^39, 40^ The few oocytes produced in *skr-1/2* RNAi-treated germlines lack chiasmata,^16^ suggesting a requirement for SCF complexes in synapsis and crossover formation. However, the role of SCF during meiotic prophase has not been established in *C. elegans*.

Here we demonstrate that SKR-1 and SKR-2 serve as structural components of the SC, independent of their canonical roles within the SCF complex. The recruitment of SKR-1/2 to the SC requires their dimerization through a conserved hydrophobic interface that overlaps with the binding sites for CUL-1 and F-box proteins, providing the molecular basis for their dual function. Remarkably, SKR-1 interacts with the previously known SC proteins and forms a soluble complex *in vitro.* Our findings reveal how a highly conserved cell cycle regulator is co-opted to interact with rapidly evolving proteins to construct an essential meiotic scaffold.

## Results

### Two paralogous Skp1-related proteins in *C. elegans*, SKR-1 and SKR-2, associate with SC components

The *skr-1* and *skr-2* genes are adjacent with one another within a 2.7-kb region on Chromosome I. The Basic Local Alignment Search Tool (BLAST) using SKR-1/2 as queries finds a single high-confidence hit in related *Caenorhabditis* species, which are present in syntenic regions with the identical set of neighboring genes as the *C. elegans skr-1/2* (**Figure 1B**). Therefore, *skr-1* and *skr-2* are likely born from a recent duplication event within the *C. elegans* lineage.

To gain insight into the function of SKR-1/2 during meiotic prophase, we used CRISPR-mediated genome editing to insert a 3xFlag epitope into the C-terminus of SKR 2 **(Figure S1)**. We then immunoprecipitated SKR-2::3xFlag from isolated germline nuclei and analyzed the proteins associated with SKR-2 by mass spectrometry (**Figure 1C and Table S1**). As expected, components of the SCF complex that directly bind Skp1, such as CUL-1 and germline-enriched F-box proteins,^18, 41^ and subunits of the 20S proteasome were specifically purified with SKR-2 (**Figure 1D**). Remarkably, all six previously known SC proteins (SYP-1, -2, -3, -4, -5, and -6) were also purified with SKR-2 and among the most abundant proteins in the SKR-2 immunoprecipitates. This is consistent with the recent evidence that SKR-1 and SKR-2 are found in the immunoprecipitates of SC components^42, 43^, supporting the conclusion that they associate with the SC in *C. elegans*.

### SKR-1 and SKR-2 localize to the SC central region

We next inserted an HA tag at the N-terminus of SKR-1 using CRISPR to determine its localization **(Figure S1)**. Immunofluorescence revealed that SKR-1 appears as bright punta during early meiotic prophase, which colocalize with SC protein aggregates known as polycomplexes **(Figure S2A)**. In pachytene, SKR-1 was found along chromosomes and mirrored the localization of SYP-5. During diplotene, when the SC central region disassembles, SKR-1 was dispersed into the nucleoplasm, but still enriched on the SC “short arm” and in the nucleolus **(Figure S2B)**. SKR-2 showed a similar pattern of localization and was detected along the SC in pachytene (**Figure 2A**). In mutants lacking an axis component HIM-3, SC assembly is severely disrupted, and SC proteins form either short stretches or nuclear aggregates,^39, 44^ where SKR-2 was also found (**Figure 2A**). Furthermore, in *syp-2* mutants lacking the SC,^39^ SKR-2 was no longer detected on meiotic chromosomes. Thus, SKR-1 and SKR-2 associate with the SC central region and have characteristics similar to other SC proteins.^42, 43, 45, 46^

**Figure 2.**
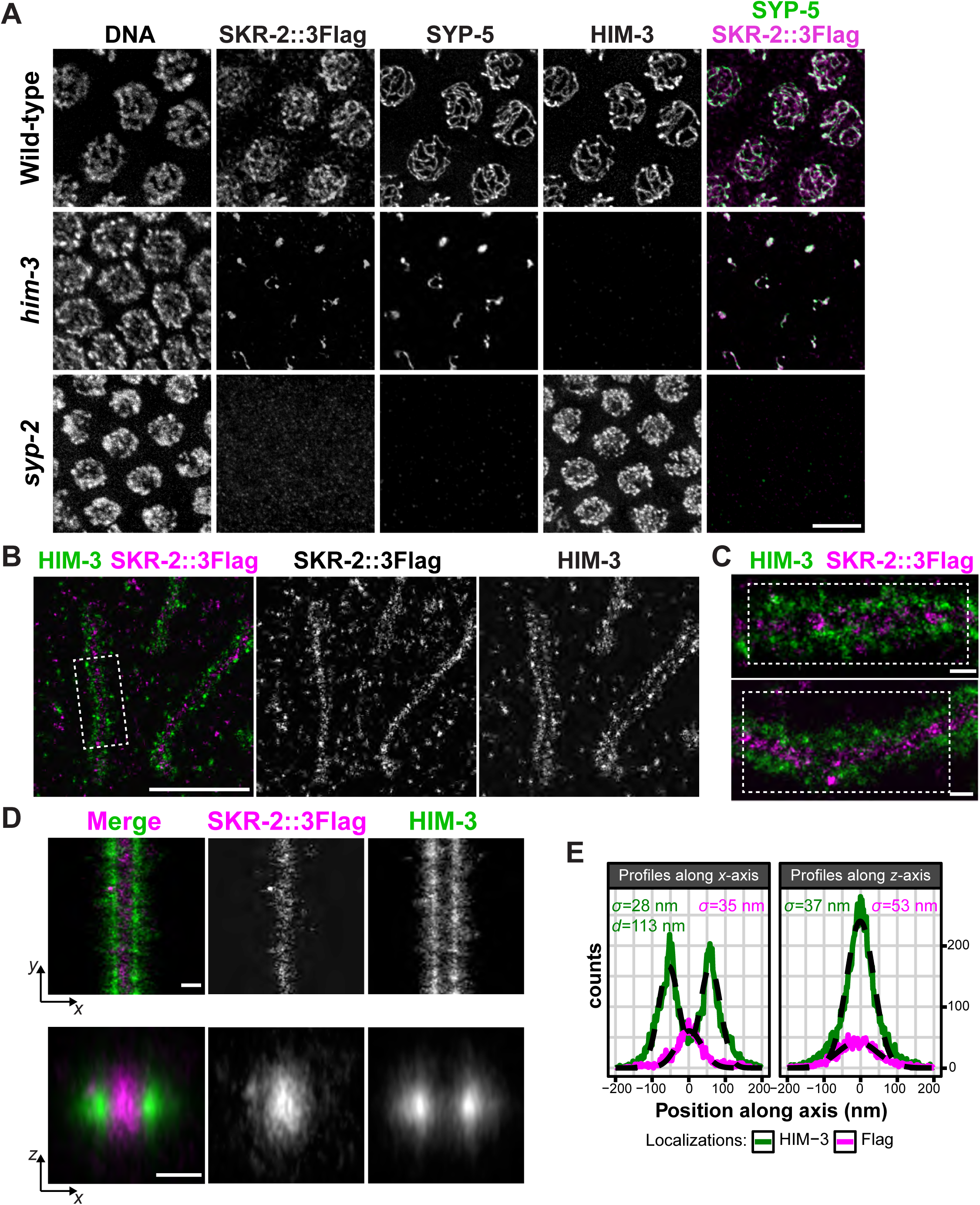
SKR-2 localizes to the SC central region. (A) Pachytene nuclei from worm strains expressing SKR-2::3Flag in the wild-type, *him 3(gk149)*, and *syp-2(ok307)* backgrounds were stained for DNA, Flag, SYP-5, and HIM-3. Scale bar, 5 µm. (B) Single-molecule localization microscopy (SMLM) images (Fourier ring correlation (FRC) resolution = 35 nm) showing a late pachytene nucleus stained for HIM-3 (green) and a Flag epitope tag fused to the C-terminus of SKR-2 (magenta). Scale bar, 1 µm. (C) Zoomed-in view of two stretches showing HIM-3 (green) and SKR-2::Flag (magenta) in late pachytene reveals the localization of SKR-2 within the center of the SC (FRC resolution = 40 nm). Scale bars, 100 nm. (D) SMLM localizations within the three boxed regions in (B) and (C) were straightened, aligned, and rendered to create averaged frontal (*xy*) and cross-sectional (*xz*) views. Scale bars, 100 nm. (E) To determine the distribution of HIM-3 and SKR-2 within the SC, histogram counts (2 nm wide bins) of straightened and aligned SMLM localizations along the *x* and *z* axes were fitted with Gaussian distributions (black dashed lines). The respective standard deviations (*σ*), and peak-to-peak distance of HIM-3 (*d*) are indicated in the plot.

We also visualized both SKR-1 and SKR-2 relative to axis proteins, HTP-3 or HIM-3, using 3D single-molecule localization microscopy (SMLM).^42, 47^ The HA tag on the N terminus of SKR-1 and the 3xflag tag on the C-terminus of SKR-2 are positioned at the center of the SC between the two chromosome axes (**Figures 2B-E and S2C**). Both epitopes were confined to a narrow z plane in the cross-sectional view, indicating that SKR-1 and SKR-2 reside at the core of the *C. elegans* SC. This is in contrast to the localization of mouse SKP1 within the SC, which appears as two parallel stretches near chromosome axes on synapsed chromosomes.^24^

### SKR-1 and SKR-2 are essential for SC assembly

To determine the role of SKR-1 and SKR-2 during meiotic prophase, we first generated a null allele *skr-2(kim66),* which harbors a 50-bp deletion immediately after the start codon, resulting in a premature stop after five amino acids (**Figure 3A**). *skr-2(kim66)* animals were indistinguishable from the wild-type, displaying normal brood sizes and egg viability **(Figure S1)**. This is in contrast to the previously characterized *skr-2(ok1938)* allele, which removes the second half of its coding sequence and a promoter region upstream of *skr-1*, reducing the expression level of *skr-1* by approximately 40% at the mRNA level.^48^ *skr-2(ok1938)* mutants laid only 12 eggs on average (compared to 317 in wild-type animals), none of which survived **(Figure S1)**. The reduction in brood size in *skr-2(ok1938)* animals is largely due to a failure to exit pachytene **(Figure S3A-B)**, as previously reported in *skr-1/2* RNAi*-*treated animals.^16^ *C. elegans* has six pairs of chromosomes, and six DAPI bodies representing the six bivalents are observed in diakinesis oocytes of wild-type and *skr-2(kim66)* animals (**Figures 3B-C**). However, 11-12 univalents were consistently detected in a few oocytes of *skr-2(ok1938)* mutants that exited pachytene, indicating a failure in crossover formation. Thus, while SKR-2 is dispensable, both SKR-1 and SKR-2 contribute to cell cycle progression in the germline and meiotic crossing over.

**Figure 3.**
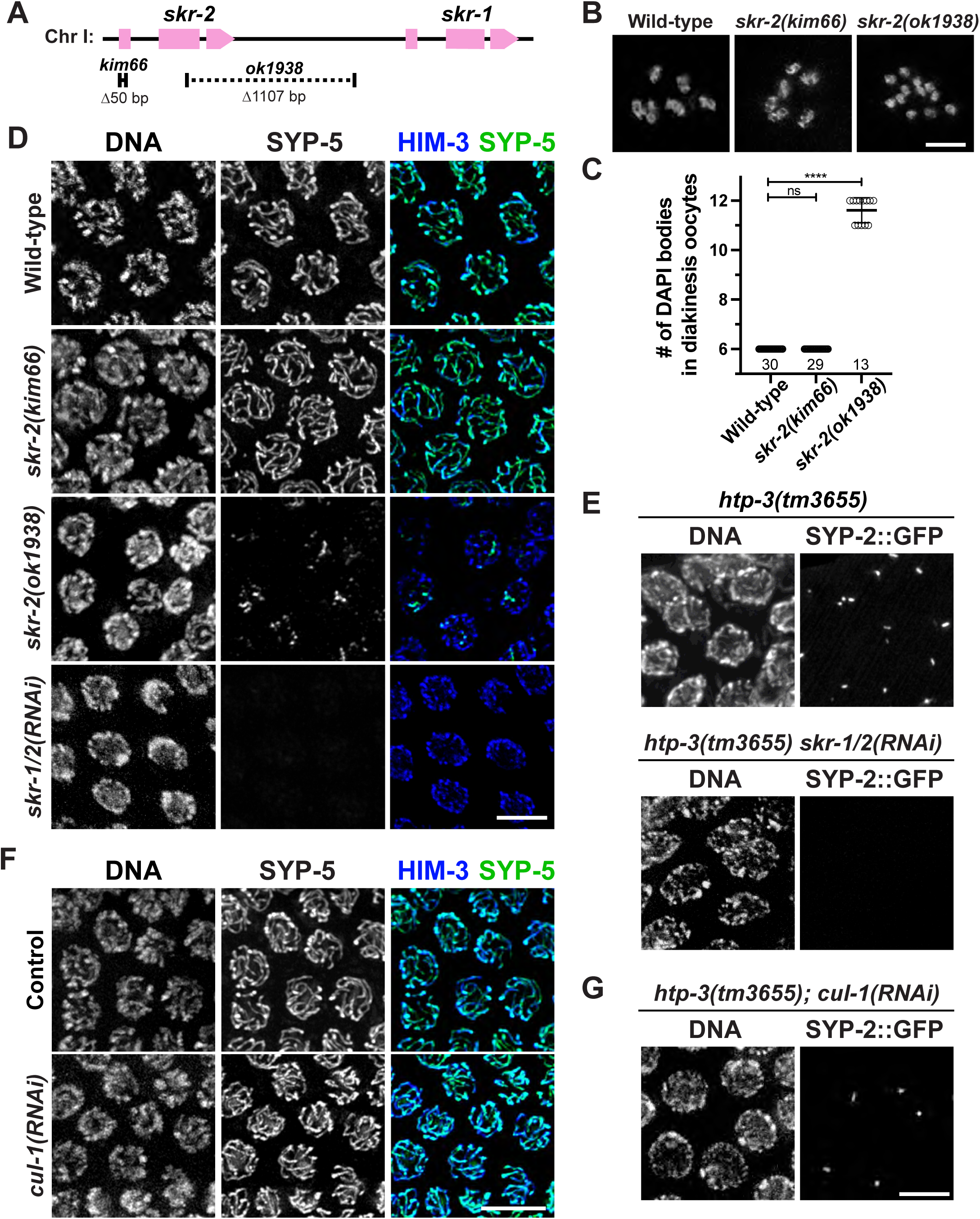
SKR-1 and SKR-2, but not CUL-1, are required for SC assembly. (A) Schematic showing the *skr* alleles used in this study. (B) Oocytes nuclei at diakinesis from indicated genotypes were stained with DAPI. Scale bar, 3 µm. (C) Graph showing the number of DAPI bodies in diakinesis oocytes. The mean ± S.D is shown. ns, not significant (p>0.9999); **** p<0.0001 by Mann-Whitney *U* test. The numbers of oocytes scored are indicated below. (D and F) Immunofluorescence images of pachytene nuclei from the indicated genotypes showing DNA, SYP-5 (green), and HIM-3 (blue) staining. Scale bars, 5 µm. (E and G) *htp-3(tm3655)* animals expressing SYP-2::GFP were treated with control (feeding), *skr-1/2* (microinjection), and *cul-1* RNAi (feeding), dissected and stained for DNA and GFP. Pachytene nuclei are shown. Scale bars, 5 µm.

As expected based on the egg viability data, *skr-2(kim66)* animals displayed normal SC assembly (**Figure 3D and S3A**). In contrast, *skr-2(ok1938)* animals showed a lack of SYP-5 signal in most pachytene nuclei, and partial SC stretches formed only in late pachytene (**Figures 3D and S3B**). Due to these severe synapsis defects, the leptotene/zygotene stage, marked by the crescent-shaped nuclear morphology, was greatly extended in *skr-2(ok1938)* animals, as previously observed with *skr-1/2* RNAi.^16^ We attempted to generate a mutant that deletes both *skr-1* and *skr-2*. However, we were not able to recover homozygous animals due to their essential function during embryo development. Thus, we turned to RNAi to knock down both *skr-1* and *skr-2* in adult animals and assessed their roles in SC assembly during meiosis. *skr-1/2* RNAi by feeding resulted in strong synapsis defects albeit to varying degrees (**Figures 3D and S3C**). To achieve a more penetrant knockdown, we microinjected *skr-1* dsRNA (1-2 µg/µl) into both gonads of L4 animals and monitored SC morphology using a worm strain expressing SYP-2::GFP. Because *skr-1* and *skr-2* share 83% nucleotide sequence identity, *skr-1* RNAi is expected to also reduce *skr-2* expression. By 2-3 days post-injection, 22 out 24 injected hermaphrodites displayed a complete loss of SYP-2::GFP signal in the germlines. SC polycomplexes were also eliminated when *skr-1* dsRNA was injected into animals lacking an axis component HTP-3 (**Figure 3E**). Thus, we concluded that SKR-1 and SKR-2 are required for SC assembly.

### CUL-1 does not localize to the SC and is dispensable for synapsis

Next, we tagged CUL-1 with the small epitope Ollas at the C-terminus to characterize its localization in the *C. elegans* germline **(Figure S1)**. CUL-1 was expressed throughout the germline, but, unlike SKR-1/2, it did not show any enrichment on meiotic chromosomes **(Figures S4A-B)**. Furthermore, CUL-1 was not found in short SYP stretches in *him-3* mutants **(Figure S4B)**, demonstrating that CUL-1 does not associate with the SC. Worms homozygous for a *cul-1* deletion arrest at larval stages due to its role in cell cycle.^49^ Thus, we employed RNAi to knock down *cul-1* expression in later development. An immunoblot showed that >90% of CUL-1 was depleted by RNAi **(Figure S4C)**, which was sufficient to cause a delay in the degradation of PPM-1.D by SCF^PROM-1^ **(Figures S4D-E)** as previously reported.^34, 35^ However, SC assembly occurred normally in *cul-1* RNAi-treated animals (**Figure 3F**), and knockdown of *cul-1* did not affect SC polycomplexes in *htp-3* mutants (**Figure 3G**). Thus, CUL-1 appears to be dispensable for synapsis, suggesting that SKR-1 and SKR-2 promote SC assembly outside the context of SCF complexes.

### SKR-1 dimerizes through a conserved hydrophobic interface that overlaps with the core F-box protein-binding interface

Biochemical analyses of recombinant Skp1 from guinea pigs and *Dictyostelium* have revealed that it forms a stable homodimer at higher concentrations.^50–52^ Given the bilateral symmetry of the SC, we hypothesized that SKR proteins might also dimerize within it during *C. elegans* meiosis. This is supported by the detection of peptides unique to SKR-1 in our SKR-2 immunoprecipitates (**Figure 1D**x), suggesting that SKR-1 and SKR-2 may form a heterodimer *in vivo*. To determine the oligomeric state of SKR proteins, we expressed and purified 6His-tagged SKR-1 from *E. coli* (**Figure 4A**) and measured its absolute molar mass using size-exclusion chromatography and multi-angle light scattering (SEC-MALS). Recombinant SKR-1 was eluted as a single peak with a molecular weight of 40.5 ± 0.9 kDa (n=3), demonstrating that SKR-1 indeed exists as a dimer in solution (**Figure 4B**).

**Figure 4.**
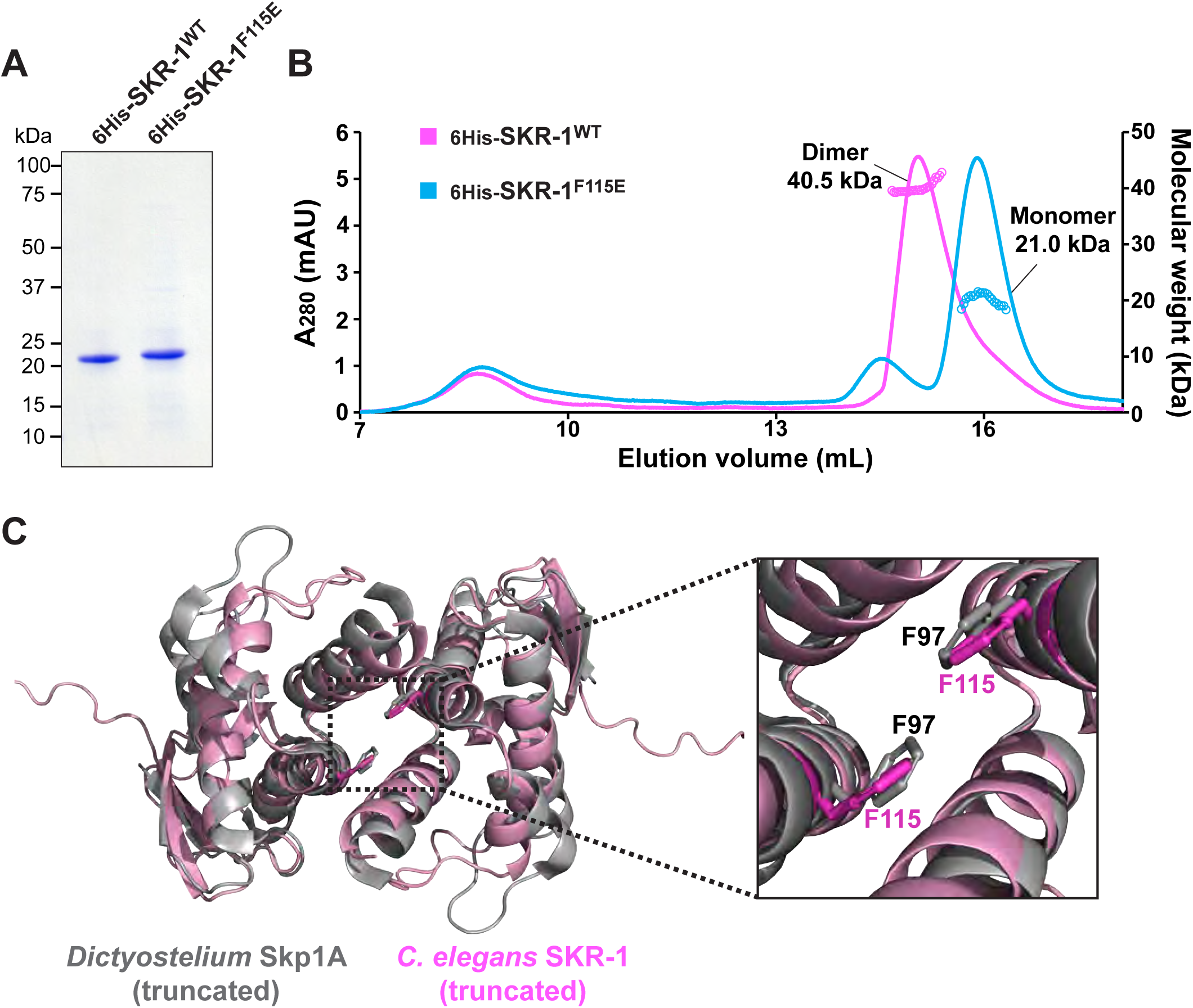
Recombinant SKR-1 exists as a dimer in solution. (A) An SDS-PAGE gel showing the purification of recombinant 6His-SKR-1^WT^ and 6His-SKR-1^F115E^. (B) SEC-MALS traces of 6His-SKR-1^WT^ (magenta) and 6His-SKR-1^F115E^ (light blue). Absorbance at 280 nm is shown on the left axis, and the measured molecular weight is shown on the right axis. (C) An AlphaFold model of *C. elegans* SKR-1 (magenta) superimposed onto the NMR structure of a truncated *Dictyostelium discoideum* Skp1A dimer (grey; PDB, 6V88).^50^ The inset shows the conserved phenylalanine (F97 in *Dictyostelium* Skp1A and F115 in *C. elegans* SKR-1) at the dimerization interface.

The dimer interface of Skp1 is conserved throughout evolution and primarily composed of a hydrophobic surface that overlaps with the core binding site for F-box domain proteins.^41, 50^ An AlphaFold model^53^ of SKR-1 superimposes well onto the NMR structure of the truncated *Dictyostelium* Skp1 dimer^50^ (**Figure 4C**). The conserved phenylalanine (F115) at the SKR-1 dimer interface overlays with F97 in *Dictyostelium* Skp1 whose substitution to glutamic acid was shown to disrupt Skp1 dimerization without affecting the binding to F-box proteins.^50^ Consistent with this, the F115E mutant of SKR-1 was monomeric in solution, as indicated by its measured molar mass of 21.0 ± 1.1 kDa (n=3) by SEC-MALS (**Figures 4A-B**).

### The dimer interface of SKR proteins is essential for SC loading and assembly

To investigate the significance of SKR dimerization in SC assembly, we introduced the F115E mutation in the *ha::skr-1* worm strain using CRISPR. The *ha::skr-1^F115E^* strain had a normal brood size and produced mostly viable progeny **(Figures S5A-B)**. Strikingly, HA::SKR-1^F115E^ protein failed to localize along the SC (**Figure 5A**), although it is expressed, albeit at a reduced level compared to the wild-type protein (**Figure 5B**). When the *skr-1^F115E^* mutation was introduced into the *skr-2(kim66)* background, a significant reduction in egg viability was observed, with only 2% of eggs surviving and 19% of the progeny being male **(Figures S5B)**. Surprisingly, SC assembly failed completely in these mutants, resulting in an extension of the transition zone corresponding to the prolonged leptotene/zygotene stage (**Figures 5C-D**). However, unlike the phenotypes seen in *skr-2(ok1938)* mutants or after *skr-1/2* RNAi,^34, 35^ there was no delay in the degradation of PPM-1.D or the onset of meiotic prophase in these animals **(Figures S5C-D)**. Additionally, meiotic cells successfully progressed through pachytene, and oocytes were produced (**Figure 5C**), indicating that the major cell cycle transitions controlled by SCF occurred normally in the germline of *skr-1^F115E^ skr-2(kim66)* mutants. Oocytes in these animals consistently displayed 10-12 DAPI-staining bodies (**Figures 5E-F**), indicating an inability to form crossovers because of synapsis failure. Together, our data demonstrate that the dimer interface of SKR proteins is essential for SC loading and assembly.

**Figure 5.**
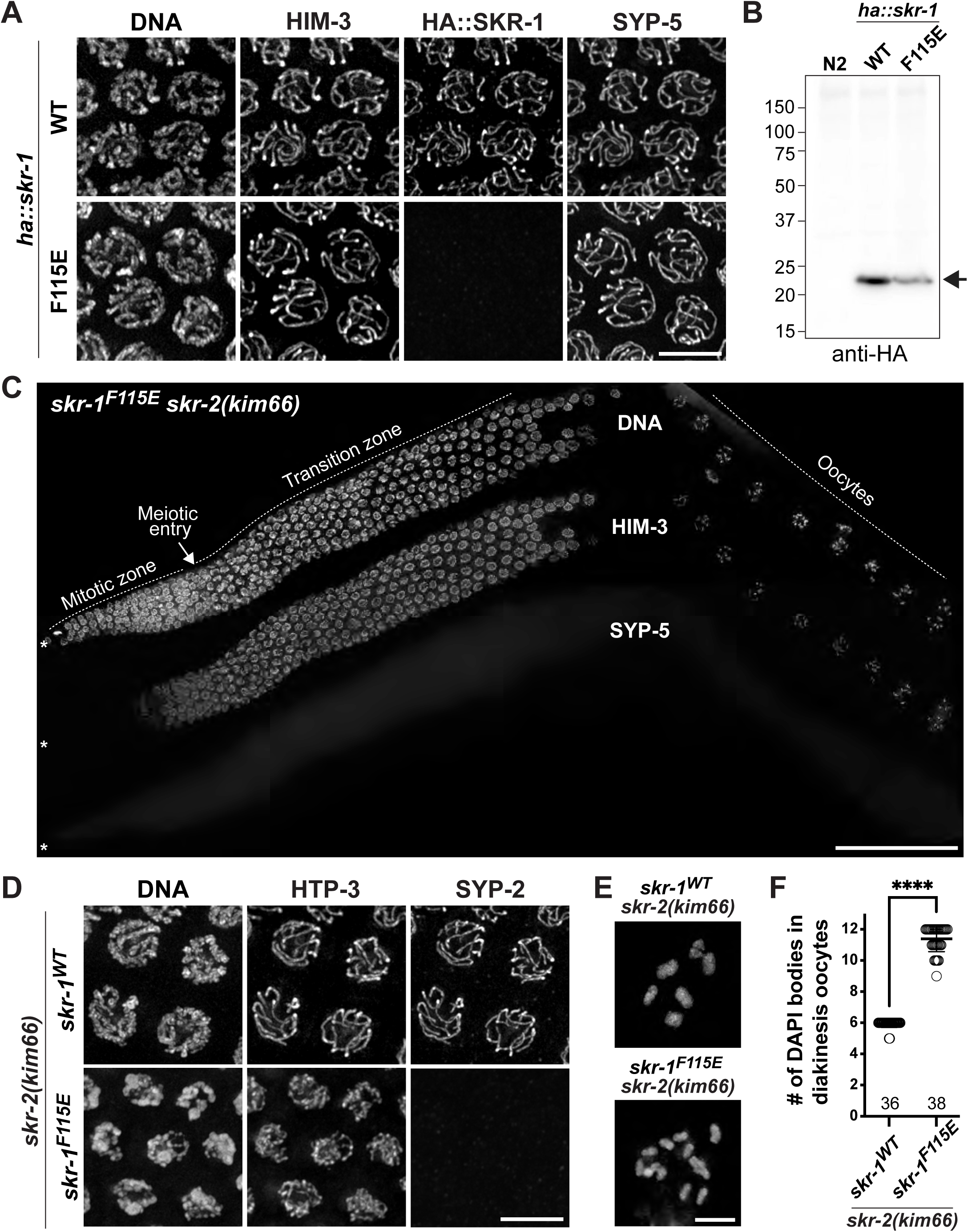
The dimerization interface of SKR proteins is essential for SC assembly. (A) Immunofluorescence images of pachytene nuclei from worm strains expressing HA::SKR-1^WT^ or HA::SKR-1^F115E^ are shown for DNA, HIM-3, HA (fused to SKR-1), and SYP-5 staining. Scale bar, 5 µm. (B) Immunoblot of worm lysates from indicated genotypes probed against the HA tag. (C) Composite immunofluorescence images of a full-length gonad dissected from *skr-2(kim66)* and *skr-1^F115E^ skr-2(kim66)* animals are shown for DNA, HIM-3, and SYP-5 staining. Asterisk (*) indicates the distal tip. Scale bar, 50 µm. (D) Immunofluorescence images of pachytene nuclei from *skr-2(kim66)* and *skr-1^F115E^ skr-2(kim66)* animals showing DNA, HTP-3, and SYP-2 staining. Scale bar, 5 µm. (E) Oocyte nuclei at diakinesis from *skr-2(kim66)* and *skr-1^F115E^ skr-2(kim66)* animals were dissected for DAPI staining. Scale bar, 3 µm. (F) Graph showing the number of DAPI bodies in diakinesis nuclei from indicated genotypes. The numbers of nuclei scored are shown below. The mean ± SD is shown; **** p<0.0001 by the Mann-Whitney *U* test.

### SKR-2 associates with the SYP proteins using the C-terminal helices

We next asked whether SKR proteins use the other SCF-forming interfaces to incorporate into the SC. The Skp1-Cul1 interaction is mediated through the BTB/POZ fold of Skp1, and mutating two residues in the H5 helix of human Skp1 has been shown to abolish its binding to Cul1.^18^ Thus, we mutated the corresponding residues in SKR-1 (N122K, Y123K) and SKR-2 (N120K, Y121K) in *C. elegans* to examine the importance of this interface in SC loading. While an *skr-2::3flag* strain harboring the N120K and Y121K mutations was fully viable **(Figure S1)**, animals homozygous for the equivalent mutations in *skr-1* could not be recovered, reflecting the essential role of SKR-1 in embryo development.^54^ Interestingly, SKR-2^N120K Y121K^ failed to localize along the SC (**Figure 6A**), even though it was clearly expressed (**Figure 6B**). SEC-MALS analyses revealed that recombinant SKR-1^N122K Y123K^ is monomeric in solution with an average molecular weight of 19.6 kDa ± 2.6 kDa (n=3) (**Figures 6C-D**), explaining the inability of SKR-2^N120K^ ^Y121K^ to load onto the SC. An AlphaFold model^53^ of the full-length SKR-1 superimposed onto the *Dictyostelium* Skp1 dimer shows that the CUL-1-binding interface of SKR proteins is within its dimer interface **(Figures S6A-B)**, suggesting that SKR dimerization is likely to interfere with CUL-1 binding.

**Figure 6.**
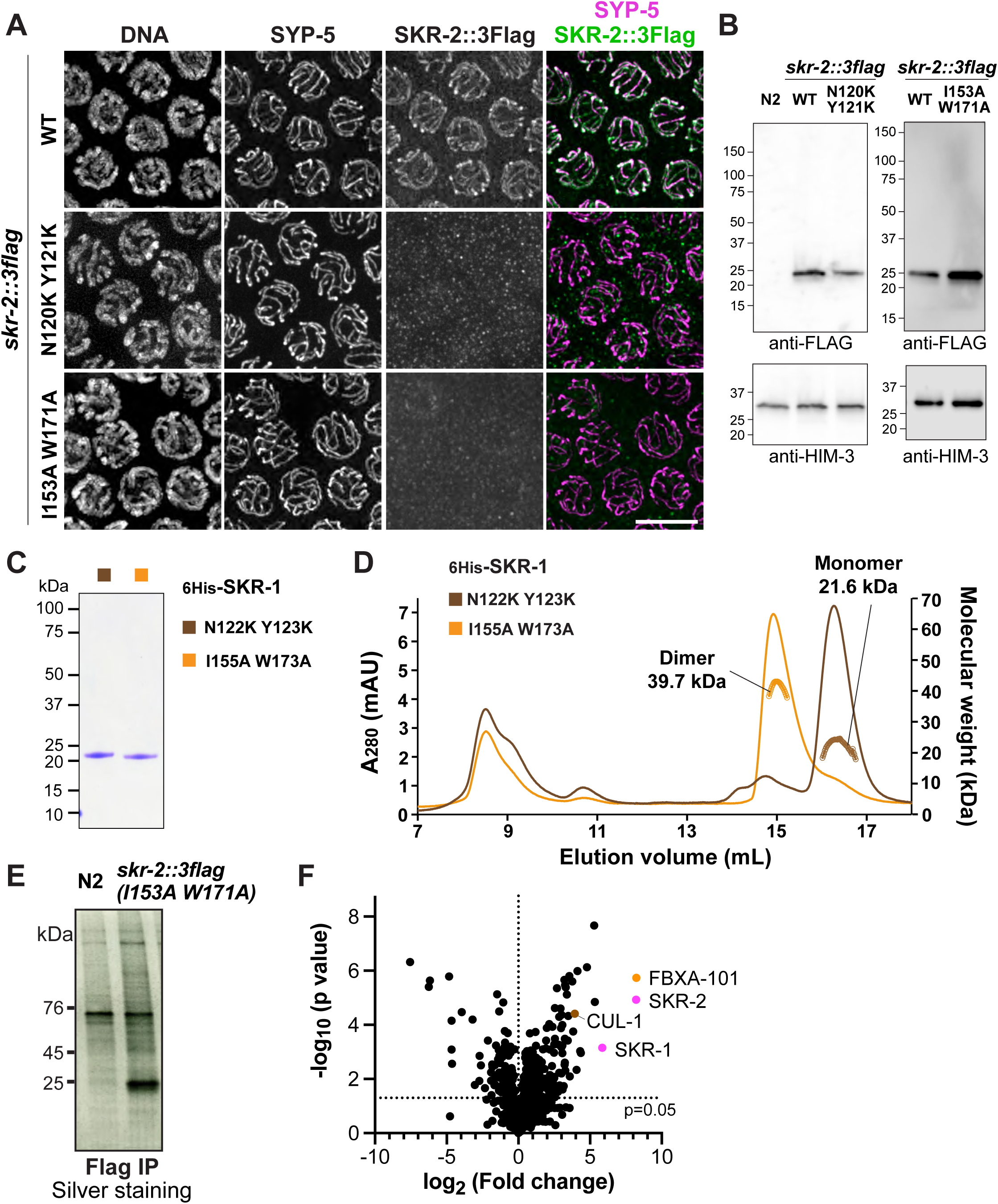
SKR-2 associates with other SC proteins using the C-terminal helices. (A) Immunofluorescence images of pachytene nuclei from *skr-2::3flag* worm strains harboring wild-type, N120K Y121K, and I153A W171A mutations. DNA, SYP-5, and SKR-2::3Flag staining are shown. Scale bar, 5 µm. (B) Immunoblots of worm lysates from the indicated genotypes probed against the FLAG tag fused to the C-terminus of SKR-2. N2 was used as a negative control, and HIM-3 was used as a loading control. (C) An SDS PAGE gel showing the purification of recombinant 6His-SKR-1^N122K^ ^Y123K^ and 6His-SKR-1^I155A^ ^W173A^. (D) SEC-MALS traces of 6His-SKR-1^N122K^ ^Y123K^ (brown) and 6His-SKR-1^I155A W173A^ (orange). The absorbance at 280 nm is shown on the left axis, and the measured molecular weight is shown on the right axis. (E) A silver-stained SDS-PAGE gel showing purified SKR-2^I153A^ ^W171A^-containing protein complexes using anti-Flag beads from worm lysates. N2 was used as a control. (F) A volcano plot showing proteins enriched in SKR-2::3flag^I153A^ ^W171A^ immunoprecipitates over N2 control. Normalized weighted spectra were transformed to logarithmic values (base 2) and analyzed by multiple unpaired t-test. The logarithm of fold change (base 2) is plotted on the x-axis, and the negative logarithm of the p-value (base 10) is plotted on the y-axis. The p-value of 0.05 is indicated by a horizontal dotted line.

Skp1 interacts with the F-box protein through a bipartite interface that involves the variable C-terminal helices of Skp1.^41^ To investigate the role of these helices in SC loading, we mutated two conserved residues in SKR-2 located at the end of H7 and H8 helices (I153A and W171A), similar to mutations that disrupt Skp1’s cell cycle function in budding yeast.^41, 55^ Importantly, these helices reside outside of the Skp1 dimer interface (Figure S6A), and recombinant SKR-1 harboring the equivalent mutations (I155A, W173A) still formed a dimer in our SEC-MALS analyses (41.0 kDa ± 1.3 kDa (n=3)) (**Figures 6C-D**). Due to the genetic redundancy between SKR-1 and SKR-2, animals harboring the *skr-2^I153A^ ^W171A^* mutation formed the SC and supported normal meiotic progression. However, SKR-2^I153A^ ^W171A^ failed to localize along the SC (**Figure 6A**), suggesting that SKR proteins require the C-terminal helices to associate with SC proteins. To test this, we immunoprecipitated 3Flag-tagged SKR-2^I153A^ ^W171A^ from isolated germline nuclei and analyzed its interacting proteins using mass spectrometry (**Figure 6E**). Indeed, SKR-2^I153A^ ^W171A^ failed to pull down the SYP proteins and most F-box proteins (**Figure 6F and Table S2**), confirming that SKR proteins interact with the other SC proteins using their C-terminal helices.

### SKR-1 forms a soluble complex with the other SC proteins *in vitro*

To demonstrate that SKR proteins are indeed structural components of the SC, we turned to *in vitro* reconstitution with recombinant proteins. Our previous attempts to co-express and purify the SYP proteins from bacteria were unsuccessful due to their insolubility. However, adding SKR-1 to the polycistronic expression vector was sufficient to solubilize all SYP proteins and allowed the purification of 6His-SYP-3 along with untagged SYP-1, SYP-2, SYP-4, SYP-5, and SKR-1 throughout multiple purification steps (**Figure 7A**). We confirmed the presence of these proteins by mass spectrometry and found that the SYP/SKR complex was eluted as a single peak from a size exclusion column (**Figure 7B**), demonstrating that SKR-1 interacts with the SYP proteins to form a homogeneous complex. These results establish SKR-1/2 as SC subunits and suggest that the entire set of SC components has now been identified in *C. elegans*.

**Figure 7.**
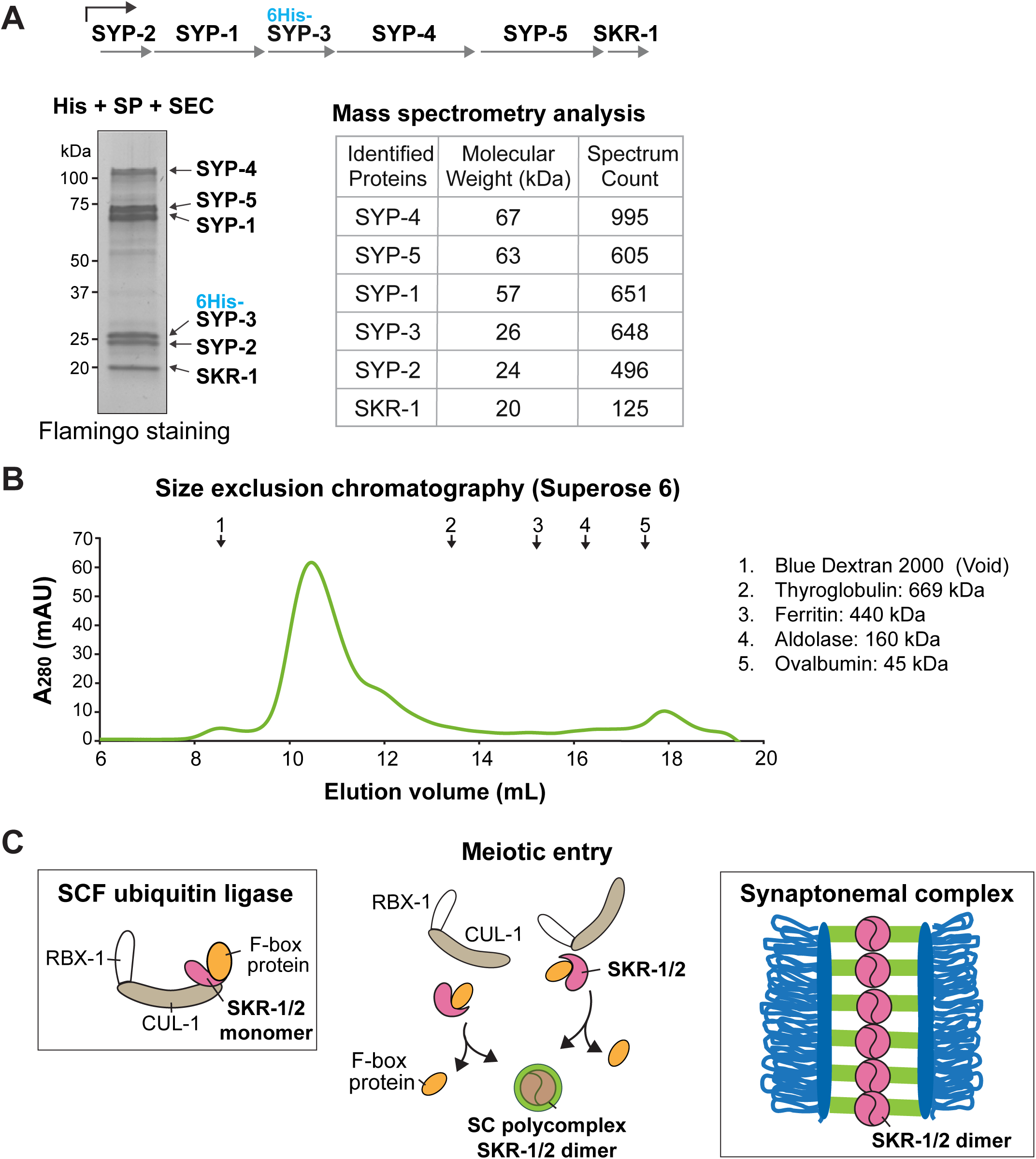
*In vitro* reconstitution establishes SKR-1/2 as structural components of the SC. (A) Schematic of the polycistronic vector used to co-express SYP proteins and SKR-1. Left: A Flamingo-stained SDS-PAGE gel of the SYP/SKR-1 complex purified using a single 6His tag on SYP-3 (His), cation exchange chromatography (SP), and size exclusion chromatography (SEC). Right: Table showing the results from mass spectrometry analysis. (B) The elution profile of the SYP/SKR-1^WT^ complex from Superose 6 (green). Arrows on the top indicate the elution volume of protein standards listed on the right. (C) Model for the moonlighting functions of SKR-1/2 within the SC. In *C. elegans*, SKR-1 and SKR-2 act as primary Skp1 adaptors for F-box proteins in the SCF ubiquitin ligase complex. SCF promotes meiotic differentiation by targeting proteins involved in the mitotic cycle for degradation.^34^ The reduced substrate availability, regulation of the binding affinity between SKR-1/2 and F-box proteins by meiotic signals, and high local concentration of SKR-1/2 may facilitate the dimerization of SKR-1/2 during meiotic entry. SKR-1/2 dimers interact with other SC proteins through the SCF-forming interfaces, leading to the formation of SC polycomplexes and synapsis between two homologous chromosomes. Two identical binding interfaces generated by the dimerization of SKR-1/2 may underlie the head-to-head symmetry of the SC.

## Discussion

Here, we demonstrate that SKR-1 and SKR-2 promote synapsis independently of their role as part of the SCF ubiquitin ligase. SC assembly requires the dimerization of SKR-1/2 through an interface that overlaps with the binding sites for Cul1 and F-box proteins.^18, 41, 50^ Therefore, loading of SKR-1/2 to the SC and SCF formation are mutually exclusive, providing the molecular basis for their dual function (**Figure 7C**). Using a separation-of-function mutation that disrupts Skp1 dimerization without impairing SCF activity, we have shown that worms carrying these mutations display phenotypes indistinguishable from those lacking other SC proteins.^39, 40, 42, 43, 45, 46^ Furthermore, a soluble protein complex can be reconstituted with SKR-1 and the five SYP proteins *in vitro*, which likely represents the basic building block for SC assembly. Our findings establish SKR-1/2 as *bona fide* subunits of the *C. elegans* SC.

Although the organizing principles of the *C. elegans* SC are currently unknown, we speculate that the dimerization of SKR proteins generates two identical interfaces for interacting with other SC proteins and contributes to its head-to-head symmetry. This mechanism might be similar to the tetramerization of SYCP1’s N-terminal region at the core of the mammalian SC, which recruits central element proteins and bifurcates into two elongated C-terminal coiled-coils toward chromosome axes.^56, 57^ While proteins that make up the SC are known for their rapid evolution and prevalent coiled-coil domains,^58, 59^ Skp1 proteins do not share these characteristics and exhibit a high degree of sequence conservation across eukaryotes.^60^ It is yet to be determined how widespread the structural function of Skp1 is conserved beyond *C. elegans* and their evolutionary history. Nevertheless, our study has uncovered a fascinating example of repurposing a highly conserved cell cycle regulator as a component of an essential meiotic scaffold.

The SCF ubiquitin ligase is active in the *C. elegans* germline and controls major cell cycle transitions throughout meiotic progression.^16, 34, 37, 38^ This raises the question of how SKR proteins switch between their canonical role as part of the SCF ubiquitin ligase and the structural function within the SC. Just before meiotic entry, an F-box protein, PROM-1, is synthesized and forms SCF^PROM-1^ complexes to downregulate proteins involved in the mitotic cell cycle.^34–36^ Since substrate availability enhances the stability of its cognate SCF complex,^61, 62^ degradation of SCF^PROM-1^ substrates might promote the rapid exchange of PROM-1-SKR-1/2 modules from CUL-1, leading to eventual disassembly of SCF^PROM-1^ complexes.

The dissociation constant (*K*_D_) for Skp1 dimerization ranges from 1 to 2.5 µM,^50, 51^ while Skp1’s binding affinity for F-box proteins is approximately 100-fold higher (*K*_D_ ∼25 nM).^63^ Thus, the dimerization of SKR-1/2 is expected when the local concentration of SKR-1/2 rises above the micromolar concentration. Although the endogenous concentration of SKR-1/2 in the *C. elegans* germline is unknown, the concentration of Skp1 in mammalian cells was measured to be within the range of 1-2 µM,^61^ which can permit Skp1 dimerization. Condensation of the SC materials into phase-separated polycomplexes may increase the local concentrations of SKR-1/2 and SYP proteins,^64^ enabling the dimerization of SKR-1/2 and the formation of building blocks required for SC polymerization (**Figure 7C**). We also speculate that signals produced during meiotic entry might weaken the interaction between SKR-1/2 and F-box proteins and help exclude F-box proteins from phase-separated SC compartments. Intriguingly, two conserved residues in the C-terminal helices of SKR-1 have been mapped to be phosphorylated *in vivo*, and phosphorylation at the equivalent residue in budding yeast Skp1 has been shown to impair its interaction with an F-box protein, Met30.^65^ Therefore, we envision a scenario in which meiotic kinases promote the dimerization of SKR-1/2 and condensation of SC proteins, thereby coupling SC assembly with the onset of meiotic differentiation.

The structural role of SKR proteins within the *C. elegans* SC is analogous to the function of Skp1 at the budding yeast centromere. Skp1 forms the CBF3 complex with three kinetochore proteins, Ctf13, Ndc10, and Cep3, which bind the centromeric DNA and direct the assembly of centromeric nucleosomes.^55, 66^ Similarly to the *C. elegans* SC, Skp1 is the only component that has clear orthologs in higher eukaryotes, while the other three proteins are poorly conserved and present only in fungi with point centromeres.^67^ Skp1 interacts with Cep3 using the Cul1-binding interface,^68, 69^ which prevents it from forming SCF ubiquitin ligases once incorporated into the CBF3 complex. Thus, Skp1 proteins have been co-opted to interact with different sets of proteins using the SCF-forming interfaces, serving as structural components of chromosome scaffolds that are essential for both mitosis and meiosis.

The formation of the CBF3 complex is known to significantly enhance the stability of Ctf13, an F-box protein, which is otherwise susceptible to degradation through SCF mediated polyubiquitination.^70, 71^ It is possible that, like Ctf13, SYP proteins are protected from degradation once incorporated into the SC, although it is unknown whether SYP proteins are targeted by the SCF complex for proteolysis at the end of meiotic prophase. Regardless, the structural function of SKR-1/2 within the SC has implications for its disassembly. As the SC starts to disassemble at the end of pachytene, germline-enriched F-box proteins could potentially function as a “molecular sponge” by sequestering SKR 1/2, thereby preventing their dimerization and accelerating desynapsis. Given the high affinity between SKR-1/2 and F-box proteins, the mere presence of F-box proteins will be sufficient to outcompete for SKR-1/2 binding and may not require the SCF ubiquitin ligase activity.

Our work in *C. elegans* has uncovered a remarkable case of meiotic regulation, in which highly conserved Skp1 proteins have acquired a novel function as structural components of the SC, outside the context of SCF ubiquitin ligases. Future work identifying the regulatory mechanisms that control the switch between their dual roles and delineating the protein-protein interactions between SKR-1/2 and the other SC proteins will provide crucial insights into the organizing principles of the SC and how its assembly and disassembly are coupled to cell cycle progression.

## Methods

### Synteny analysis of *skr-1* and *skr-2*

SKR-1 orthologs were identified by using *C. elegans* SKR-1 and SKR-2 protein sequences to query the proteomes of 19 *Caenorhabditis* species using BLASTP^72^ implemented in WormBase or Wormbase Parasite databases. Most queries produced a high-confidence hit. The syntenic locus of each identified *skr-1/2* ortholog was annotated according to orthologous genes in *C. elegans*.

### *C. elegans* strains and CRISPR-mediated genome editing

All strains used in this study were maintained on Nematode growth medium (NGM) plates seeded with OP50-1 bacteria under standard conditions.^73^ All experiments were performed at 20°C. All *C. elegans* strains were derived from Bristol N2 background. **Tables S3** and **S5** summarize all mutations and strains used in this study.

To generate strains expressing HA::SKR-1 and SKR-2::3Flag and a null allele *(kim66)* of *skr-2*, N2 or worms expressing GFP::COSA-1 were injected with 0.25 µg/µl of Cas9 (2 nmol) complexed with 10 µM tracrRNA/crRNA oligos (IDT), 40 ng/µl of pRF4::rol-6(su1006), and a ssDNA oligo (200 ng/µl) (IDT) with 35 bp homology arms on both sides as a repair template **(Table S4)**. F1 progeny were lysed and genotyped by PCR to detect successful edits. The correct insertion was validated by Sanger sequencing.

To generate the F115E mutation of *skr-1* (*kim74*) and N120K Y121K *(kim76)* or I153A W173A mutations (*kim78)* of *skr-2*, YKM616 (*ha::skr-1*) or YKM219 (*skr-2::3flag*) worms were injected respectively with 16 µM of Cas9 complexed with 16 µM tracrRNA/crRNA oligos (IDT), 5 ng/µl of pCFJ104, 2.5 ng/µl of pCFJ90, and a ssDNA oligo (100 ng/µl) (IDT) as a repair template **(Table S4)**. F1 progeny were lysed and genotyped by PCR to detect successful edits. The correct insertion was validated by Sanger sequencing.

The worm strain expressing SYP-2::TEV::GFP (YKM853) was made by InVivo Biosystems using CRISPR.

### Immunoprecipitation and mass spectrometry analysis of SKR-2-containing protein complexes

N2, YKM219 (*skr-2::3flag*), and YKM942 (*skr-2::3flag^I153A^ ^W171A^)* worms were grown in liquid culture in a developmentally synchronous manner at 20°C. Young adult worms were harvested by sucrose flotation, frozen in liquid nitrogen, and ground into a frozen powder using a Retsch Mixer Mill. The frozen powder was then resuspended in 2X volume of nuclear purification buffer (50 mM HEPES pH, 7.5, 40 mM NaCl, 90 mM KCl, 2 mM EDTA, 0.5 mM EGTA, 0.2 mM DTT, 0.5 mM PMSF, 5 mM spermidine, 0.247 mM spermine, cOmplete Protease Inhibitor Cocktail (Sigma 11697498001)) and was passed through two 40 µm, two 30 µm, and two 20 µm cell strainers (pluriSelect 43-50040-50, 43-50030-50, 43-50020-50) to isolate germline nuclei as previously described.^74^ The nuclei were collected by centrifugation at 3100 rpm for 6 min at 4°C, and the pelleted nuclei were resuspended in 2X the volume of lysis buffer (50 mM Tris-HCl pH, 8.0, 150 mM NaCl, 5 mM MgCl_2_, 1 mM EGTA, 0.1% NP-40, 10% glycerol, cOmplete Protease Inhibitor Cocktail). The isolated nuclei were transferred to a new tube and nutated for 10 min at 4°C for lysis to complete. The nuclear lysates were cleared by centrifugation at 4000 rpm for 10 min, and the supernatant was incubated with anti-FLAG M2 magnetic beads (Sigma M8823) for 10-20 min at 4°C to prevent the exchange of SCF subunits post-cell lysis.^61^ The beads were then washed three times in the lysis buffer supplemented with 300 mM NaCl, and SKR-2-containing protein complexes were eluted overnight in 100 µl of the lysis buffer containing 2 mg/mL of 3xFlag peptide (Sigma F4799). Eluted protein samples were digested with trypsin and analyzed by MudPIT in the Mass Spectrometry and Proteomics Facility at Johns Hopkins School of Medicine.

Flag immunoprecipitation was performed in three biological replicates. Spectrum counts were normalized in Scaffold 5 (Proteome Software) by multiplying the average number of spectra across all biological samples over the total number of spectra in each sample. Normalized spectra were converted to logarithms (base 2) and were analyzed by multiple unpaired t-test (GraphPad Prism 9) to identify proteins enriched in SKR-2::3Flag IP.

### Egg count

L4 hermaphrodites were picked onto individual NGM plates and transferred to new plates every 12 hr for a total of 4-5 days. Eggs and hatched L1s were counted immediately after the transfer, and surviving progeny and males on each plate were counted when F_1_ reached adulthood.

### RNAi interference

The study used two methods to carry out RNA interference: feeding and microinjection. For feeding RNAi, N2, YKM874 (*htp-3(tm3655); syp-2::tev::gfp*), YKM218 (*cul-1::Ollas*), or YKM183 (*ppm-1.D::ha*) hermaphrodites were fed with the *Escherichia coli* strain HT115 (DE3) carrying an Ahringer RNAi library clone of *skr-1,*^75^ a Vidal RNAi library clone of *cul-1,*^76^ or the empty L4440 vector. Concentrated bacterial cultures were seeded onto RNAi plates (NGM, 1 mM IPTG, and 100 µg/mL carbenicillin), left for 1 h in the laminar fume hood to dry, and incubated for 3 days at 37°C to induce expression of dsRNA. L1 hermaphrodites were picked onto seeded RNAi plates and left for 3-4 days at 20°C before dissection for immunofluorescence. *skr-1* RNAi by feeding resulted in variable phenotypes with incomplete penetrance.

For *skr-1* RNAi by microinjection, a coding sequence within exon 2 was amplified by PCR from the genomic DNA of N2 worms using primers with the T7 promoter sequence attached to the 5’ ends (Foward, taatacgactcactataggAAACACTATCAACACTCTCC; Reverse, taatacgactcactataggATGAGCTCGAAAAGGGTTCC). Sense and antisense ssRNAs were synthesized *in vitro* using the MEGAscript T7 Transcription Kit (Thermo Fisher) and annealed by incubation at 70°C for 10 min and gradual cooling to 20°C. Upon treatment with TURBO DNAse (Invitrogen) and RNase A (Thermo Fisher), dsRNA was purified using phenol-chloroform extraction and verified by gel electrophoresis. 1-1.8 µg/µl of dsRNA were injected into both gonad arms of YKM853 (*syp-2::tev::gfp)* animals at the L4 stage. The knockdown efficiency of RNAi was optimized by monitoring the SYP-2::GFP signal.

### Immunofluorescence

Adult hermaphrodite germlines were dissected 24 hr after L4 larval stage in egg buffer (25 mM HEPES, pH 7.4, 118 mM NaCl, 48 mM KCl, 2 mM EDTA, 5 mM EGTA, 0.1% Tween-20, and 15 mM NaN_3_) except for *skr-1* RNAi-microinjected animals, which were dissected 2-3 days post-L4. Dissected germlines were fixed in 4-10 % formaldehyde before freezing in liquid nitrogen, freeze-cracked, and fixed again in -20°C methanol for 1 min. Samples were then rehydrated with PBST (13.7 mM NaCl, 0.27 mM KCl, 1 mM Na_2_HPO_4_, 0.18 mM KH_2_PO_4_, pH 7.4, 0.1 % Tween 20) and blocked with Roche blocking reagent (Sigma 11096176001) for 1 hr at room temperature. Incubation with primary antibodies was performed overnight at 4°C at the following concentrations: FLAG (mouse, 1:1000; Sigma F1804), HA (mouse, 1:500; Invitrogen 2-2.2.14), SYP-2 (rabbit, 1:500),^42^ SYP-5 (rabbit, 1:500),^42^ HIM-3 (chicken, 1:500),^42^ HTP-3 (guinea pig, 1:500),^42^ GFP booster Alexa Fluor 488 (1:200; Proteintek gb2AF488), and Ollas (rat, 1:500; Invitrogen MA5-16125). Slides were washed with PBST and incubated with secondary antibodies for 30 minutes at room temperature at a 1:200 dilution (Sigma or Invitrogen Alexa 488, Alexa 555, or Alexa 647). Slides were washed again with PBST, stained with DAPI, and mounted in ProLong Gold (Invitrogen P36930). Slides were cured for 1 day before imaging.

Most images were captured with a DeltaVision Elite system (Cytiva) equipped with a 100x oil immersion, 1.4 N.A. objective, and an sCMOS camera (PCO). 3D image stacks were collected at 0.2 µm intervals, processed by iterative deconvolution (enhanced ratio, 20 cycles), and projected using the SoftWoRx package. Composite images were assembled and colored using Adobe Photoshop. Images shown in Figure S4D were captured by a Leica Thunder Imaging System equipped with a 100x oil immersion, 1.4 N.A. objective, and a Leica K8 camera. 3D image stacks were collected at 0.2 µm intervals and processed by Large Volume Computational Clearing using Global Strategy (the feature scale: 341 nm; 92% strength; 30 iterations). Image stacks were projected using the LAS X software, and the projected images were assembled and colored using Adobe Photoshop.

### Single-molecule localization microscopy (SMLM)

#### Sample preparation

Adult worms (24 hours post-L4) were dissected and immunostained as described previously.^77^ Dissected and fixed tissue was incubated overnight at 4°C with anti-Flag (mouse, 1:200; Sigma F1804) or anti-HA (mouse, 1:200; Invitrogen 2-2.2.14), and anti-HIM-3 (rabbit polyclonal, 1:200 or 1:100; Novus Biologicals #53470002) or anti HTP-3 (chicken polyclonal, 1:200)^78^ primary antibodies. The first incubation was followed by three, 5-15 min washes with PBST to remove unbound primary antibodies. Samples were subsequently stained with F(ab’)2 fragments conjugated with organic fluorophores: AlexaFluor 647 (Thermo Fischer Scientific A37573) anti-mouse IgG (donkey polyclonal, 1:50; Jackson Immunoresearch AB_2340761), and CF680 (Biotium 92139) anti-rabbit IgG (donkey polyclonal, 1:50; Jackson Immunoresearch AB_2340586) or anti-chicken IgY (donkey polyclonal, 1:50; Jackson Immunoresearch AB_2340347). Working antibody solutions were prepared in 1x Roche Blocking Buffer (Sigma 11096176001). Samples were mounted within a custom-built holder filled with 1 ml of imaging buffer (50 mM Tris HCl, pH 8.0, 10 mM NaCl, 10% D-Glucose, 35 mM 2-mercaptoethylamine (MEA), 500 μg/mL GLOX, 40 μg/mL catalase), and sealed with parafilm.^79, 80^

#### SMLM imaging

SMLM imaging was performed at room temperature on a custom-built microscope.^81, 82^ Briefly, samples were excited by a Luxx 638 nm laser coming from a LightHub laser combiner (Omicron-Laserage Laserprodukte, Dudenhofen, Germany) and an additional booster laser (Toptica Photonics iBEAM-SMART-640-S; Gräfelfing, Germany). The two lasers were combined by a polarizing beam splitter and coupled into a square multimode fiber (Thorlabs M103L05; Newton, NJ, USA). An effective homogeneous illumination was achieved through speckle reduction by mechanical agitation of the fiber.^83^ The output of the multimode fiber was first magnified by an achromatic lens and then focused on the sample. A laser cleanup filter (390/482/563/640 HC Quad; AHF, Tübingen, Germany) was introduced in the illumination path to remove fluorescence generated by the fiber.^83, 84^ The emitted light was gathered by a high numerical-aperture (NA) oil-immersion objective (160x/1.43-NA; Leica, Wetzlar, Germany) and passed through an astigmatic lens (f = 1000 mm, Thorlabs) to allow for 3D-SMLM image reconstruction. The dual-color acquisition was achieved by ratiometrically splitting the emission light by a 665 nm long-pass dichroic mirror (Chroma, ET665lp) and by filtering the transmitted and reflected photons by a 685/70 (Chroma ET685/70m) and a 676/37 (Semrock, FF01-676/37-25) filter, respectively. Both reflected and transmitted emissions were recorded as separate channels at distinct parts of the same EMCCD camera (Photometrics, Evolve512D). During the course of SMLM acquisitions, the focus was stabilized by detecting the total internal reflection of an infrared laser (iBeam Smart, Toptica, Munich, Germany) on the coverslip using a quadrant photodiode (QPD) that was in a closed feedback loop with the piezo objective positioner (Physik Instrumente).^85^ A Field Programmable Gate Array (FPGA; Au; Embedded Micro) was used to control the lasers, switch filters and stabilize the focus.^86^ The FPGA is controlled by EMU,^87^ a custom-written plugin for Micro-Manager 2.0.0.^88, 89^ The samples were exposed to the 638 nm illumination at high irradiance of about 13 kW/cm^2^ until an appropriate fluorophore blinking rate was achieved. Subsequently, 150,000 - 200,000 frames were acquired with an exposure time of 20 ms and 20 kW/cm^2^ irradiance.^83, 84^

#### SMLM Image reconstruction and post-processing

All post-processing of raw data was performed using the SMAP software (Super-resolution Microscopy Analysis Platform, https://github.com/jries/SMAP).^90^ Single emitters were localized in the acquired SMLM images by the global fitting algorithm *globLoc*^85^ within the SMAP software using an experimental PSF model and linking all three coordinates of the emitters between the two channels. The experimental astigmatic PSF model and the transformation between the two channels were generated from *z*-stack images of fluorescent beads (TetraSpeck Microspheres, T7279, Thermo Fischer Scientific) that were acquired prior to each SMLM experiment.^91^ Subsequently, sample drift was corrected by a custom algorithm based on redundant cross-correlation within the SMAP software as described previously.^84^ Furthermore, dim and out-of-focus emitters were rejected by filtering localizations according to their localization precision and their *z* position, respectively. To this end, we have filtered all localization with *z* position outside of the -300 to 300 nm range, as well as, the localizations with localization precision above 15 nm in lateral (*xy*) and 30 nm in axial (*z*) direction. Channel assignment of the individual localizations was performed according to their relative brightness in the two channels. The super-resolved image was reconstructed by plotting Gaussians at the fitted positions with a width proportional to their localization precision and assigning the color based on the assigned channel.

#### Data analysis

To assess the quality of the SMLM images, their Fourier ring correlation (FRC) resolution^92, 93^ was estimated by the *FRC resolution* plugin from SMAP software.^90^ To create frontal and cross-sectional views of the SMLM localizations (**Figure 2 and S2**), midlines of the SC stretches were manually annotated. SMLM localizations around the annotated polyline were straightened, aligned, and rendered to create displayed views using custom MATLAB (R2022a) scripts that can be run from the SMAP software.^90^ *Line profiles* SMAP plugin was used to plot the profiles (histograms with 2 nm bin width) of straightened localizations along the *x* and *z* axes. Generated profiles for each localized SC component (HIM-3 and SKR-2::3Flag) were fitted by a double or a single peak Gaussian to describe their distributions within the annotated regions. Histogram counts and the fitted Gaussian distributions were plotted by a custom R (version 4.4.2) script using the *ggplot2* package.

### Protein expression and purification

The open reading frame of SKR-1 was synthesized as a gBlock (IDT) and inserted into the pMAL-c6T vector (NEB) to express 6His-MBP (maltose-binding protein) fused to the N-terminus of SKR-1. After confirming that SKR-1 was soluble without MBP, the coding sequence for MBP was removed by Q5 site-directed mutagenesis (NEB) to express 6His-SKR-1. The F115E, N122K Y123K, and I155A W173A mutations were introduced by Q5 site-directed mutagenesis (NEB) **(Table S6)**. To construct a polycistronic vector for expressing SYPs and SKR-1, synthetic gBlocks (IDT) or gene fragments amplified from a *C. elegans* cDNA library were ligated into pET23 (Novagen) via Gibson assembly. Each translation cassette contains the translation enhancer and Shine-Dalgarno sequence upstream of the start codon. Protein expression was induced in Rosetta2(DE3)pLysS cells (Novagen) at 15°C with 0.1 mM IPTG for 16 hours. Bacterial cell pellets were flash-frozen, resuspended in the lysis buffer (PBS, 300 mM NaCl, 50 mM imidazole, 0.5 mM EGTA, 2 mM MgCl2, and 1 mM DTT) containing cOmplete Protease Inhibitor Cocktail, and lysed by sonication. After centrifugation at 13,000 rpm (Beckman, JA-17) for 45 min, the supernatant was loaded to a HisTrap HP column (Cytiva) and washed with binding buffer (PBS, 300 mM NaCl, 50 mM imidazole, 0.5 mM EGTA, 2 mM MgCl2, and 1 mM DTT). 6His-SKR-1 proteins and the SYP/SKR-1 complex were eluted in the binding buffer with 500 mM imidazole. Eluted fractions were pooled and run over the HiTrap SP HP column (Cytiva) to remove contaminating proteins. For the variants of recombinant 6His-SKR-1, the flow-through from the HiTrap SP column was collected and centrifuged to concentrate in an Amicon Ultra-4 Centrifugal Filter (Sigma UFC8010). The SYP/SKR-1 complex eluted from the HiTrap SP column using 100 mM - 1 M NaCl gradient in 50 mM HEPES, pH 7.5, 0.5 mM EGTA, 5 mM MgCl2, 1 mM DTT was digested with trypsin and analyzed by MudPIT in the Mass Spectrometry and Proteomics Facility at Johns Hopkins School of Medicine.

An AlphaFold model^53^ of a truncated (aa 1-83, GGSG, 96-143) or the full-length *C. elegans* SKR-1 was superimposed into the NMR structure of a truncated *Dictyostelium discoideum* Skp1A homodimer (PBD 6V88)^50^ using PyMol (Schrödinger, Inc).

### Size exclusion chromatography and multi-angle light scattering (SEC-MALS)

Recombinant SKR-1 proteins (0.1 mg/mL) and the SYP/SKR-1 complex (0.1 mg/mL) were separated on a Superdex 200 10/300 GL and Superose 6 10/300 GL columns (Cytiva), respectively, that were equilibrated with 10 mM HEPES, 400 mM NaCl, 0.5 mM EGTA, 5 mM MgCl_2_, 2 mM TCEP and 5% glycerol at a flow rate of 0.1 ml/min. The light scattering and reflective index profiles of recombinant SKR-1 proteins were collected by miniDAWN and OptiLab (Wyatt Technology). Data were collected every second and analyzed by ASTRA 8 (Wyatt Technology). Bovine serum albumin (Sigma P0834) was used to calibrate the light scattering detector.

## Supplemental Figure Legends

**Figure S1 (related to Figures 1, 2, 3, and 6).**
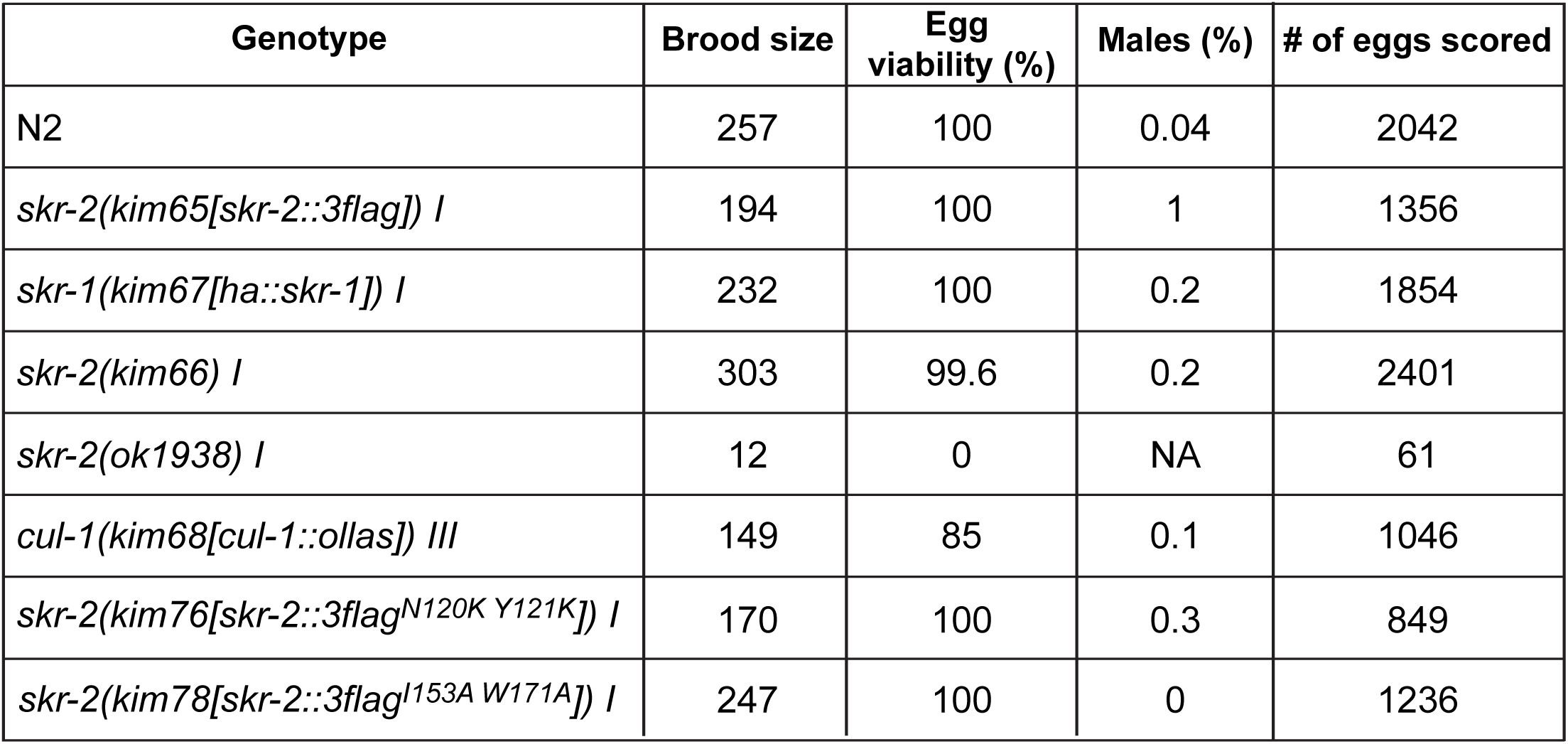
**Egg viability of *C. elegans* strains used in this study**

**Figure S2 (related to Figure 2).**
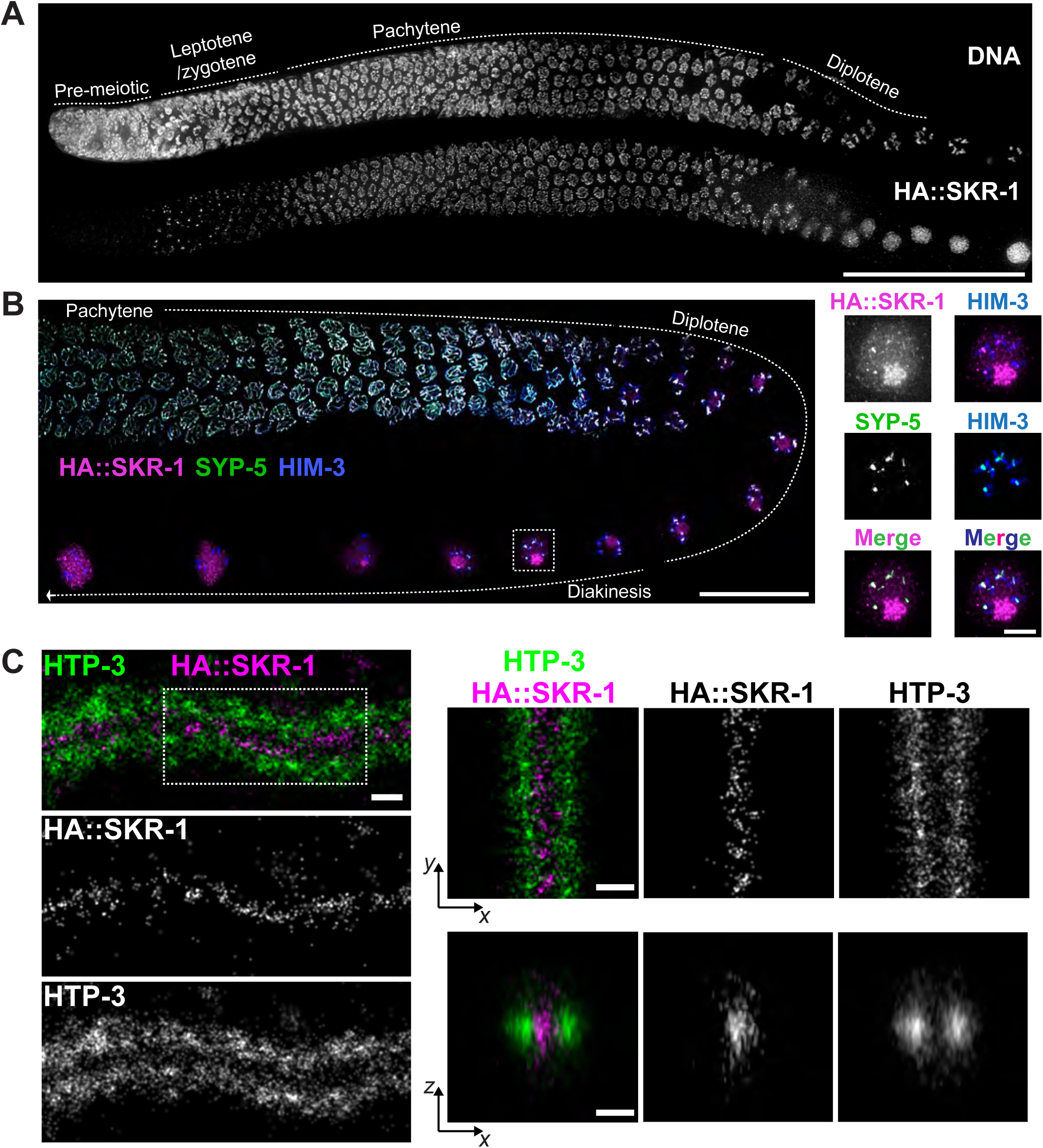
SKR-1 localizes to the SC central region. (A) A worm strain expressing HA::SKR-1 was dissected and stained for DNA and HA. Composite images are shown. Scale bar, 50 µm. (B) Composite immunofluorescence images showing HA::SKR-1 (magenta), SYP-5 (green), and HIM-3 (blue) staining in late pachytene, diplotene, and diakinesis stages. Scale bar, 20 µm. Insets: zoomed-in views of the boxed diakinesis oocyte on the left. Scale bar, 3 µm. (C) *Left:* Single-molecule localization microscopy (SMLM) images showing a stretch of a late pachytene SC stained for HTP-3 (green) and HA fused to the N-terminus of SKR-1 (magenta) (Fourier ring correlation resolution = 50 nm). *Right:* SMLM localizations within the boxed region were straightened and rendered to create frontal (*xy*) and cross-sectional (*xz*) views of the synaptonemal complex. Scale bars: 100 nm.

**Figure S3 (related to Figure 3).**
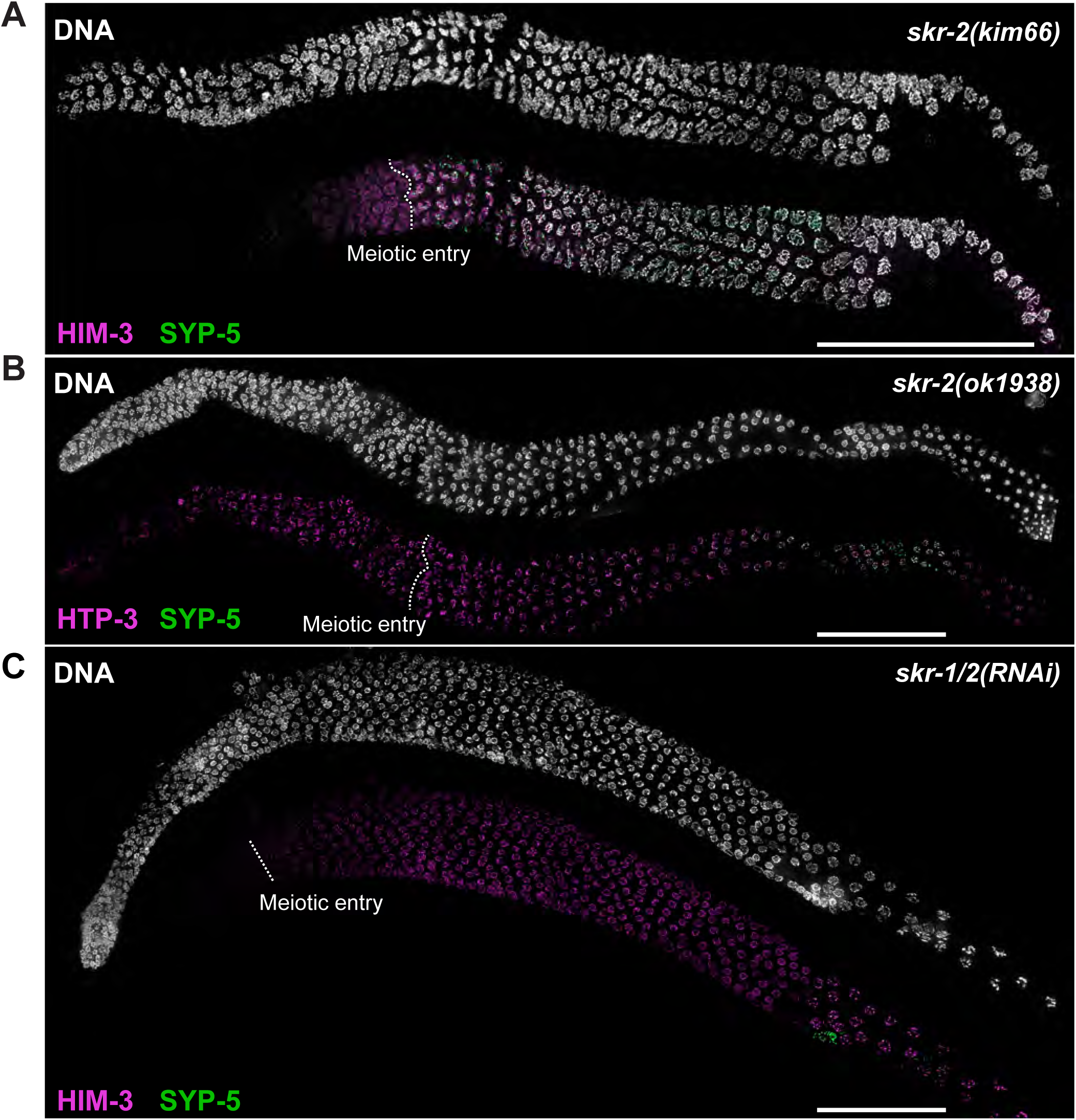
SKR-1 and SKR-2 are required for synapsis. (A-C) Composite immunofluorescence images of full-length gonads dissected from (A) *skr-2(kim66)*, (B) *skr-2(ok1938)*, and (C) *skr-1/2(RNAi)* animals are shown for DNA, SYP 5 (green), and HIM-3 or HTP-3 (magenta) staining. The onset of meiotic entry is indicated by dotted lines. The *skr-1/2 RNAi* was performed by feeding, and the effect of knockdown on pachytene exit was not fully penetrant. Scale bars, 50 µm.

**Figure S4 (related to Figure 3).**
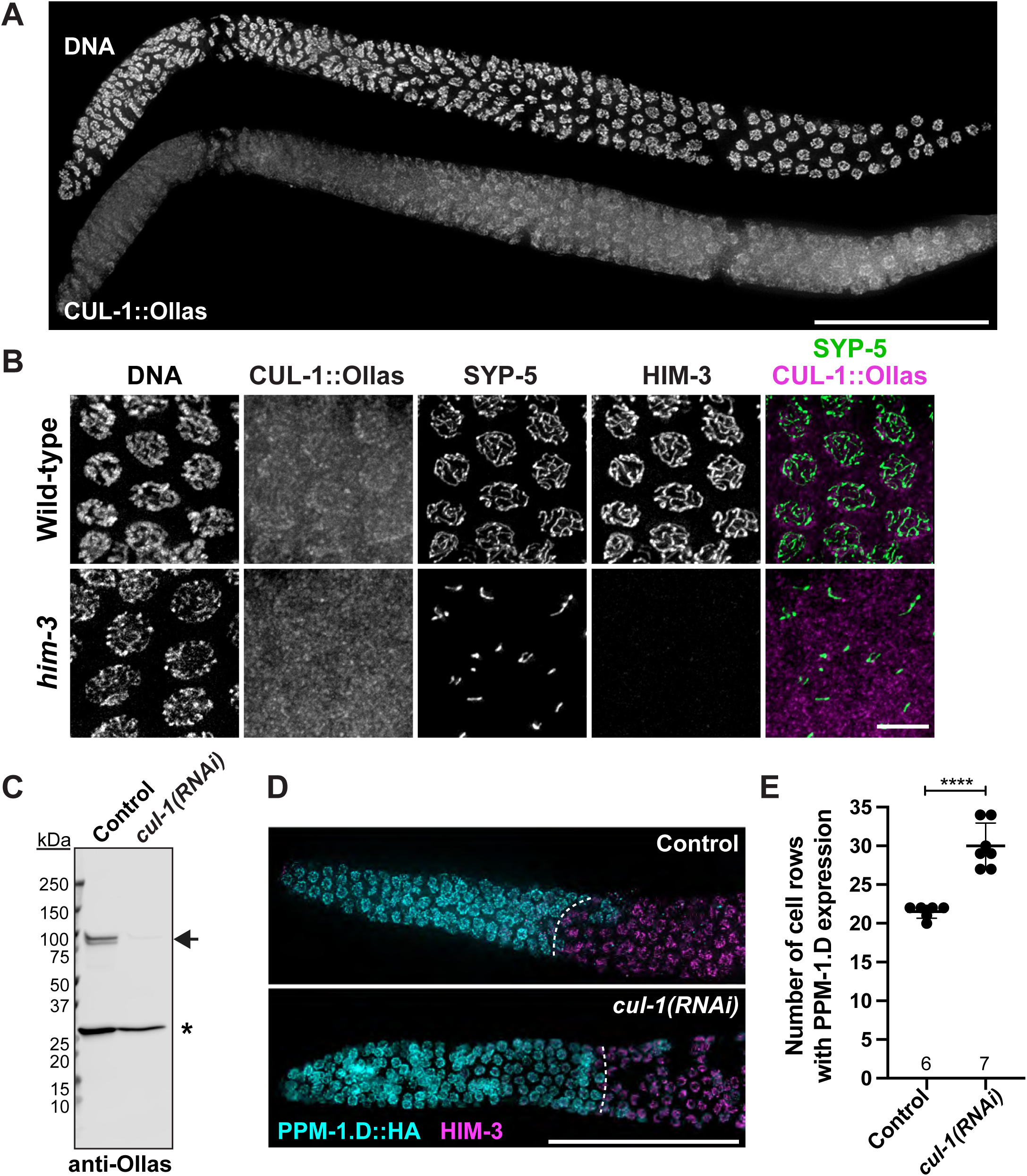
The expression of CUL-1 in the *C. elegans* germline and delayed meiotic onset upon *cul-1* RNAi. (A) Composite immunofluorescence images of a full-length gonad dissected from a worm strain expressing CUL-1::Ollas. DNA and CUL-1::Ollas staining are shown. Scale bar, 50 µm. (B) Immunofluorescence images of pachytene nuclei from wild-type and *him 3(gk149)* mutants are shown for DNA, Ollas (fused to the C-terminus of CUL-1; magenta), SYP-5 (green), and HIM-3. Scale bar, 5 µm. (C) Immunoblot of worm lysates from control and *cul-1(RNAi)* animals probed against CUL-1::Ollas (indicated by arrow). The asterisk indicates a non-specific band. (D) Composite immunofluorescence images of distal gonads dissected from control and *cul-1(RNAi)* animals stained for PPM-1.D::HA (cyan) and HIM-3 (magenta). Scale bar, 50 µm. The onset of PPM-1.D degradation is marked by white dotted lines. (E) Graph showing the number of cell rows from the distal tip that displays PPM-1.D expression. Mean ± S.D. is shown, and the numbers of gonads scored are shown below. **** p<0.0001 by Unpaired t-test.

**Figure S5 (related to Figure 5).**
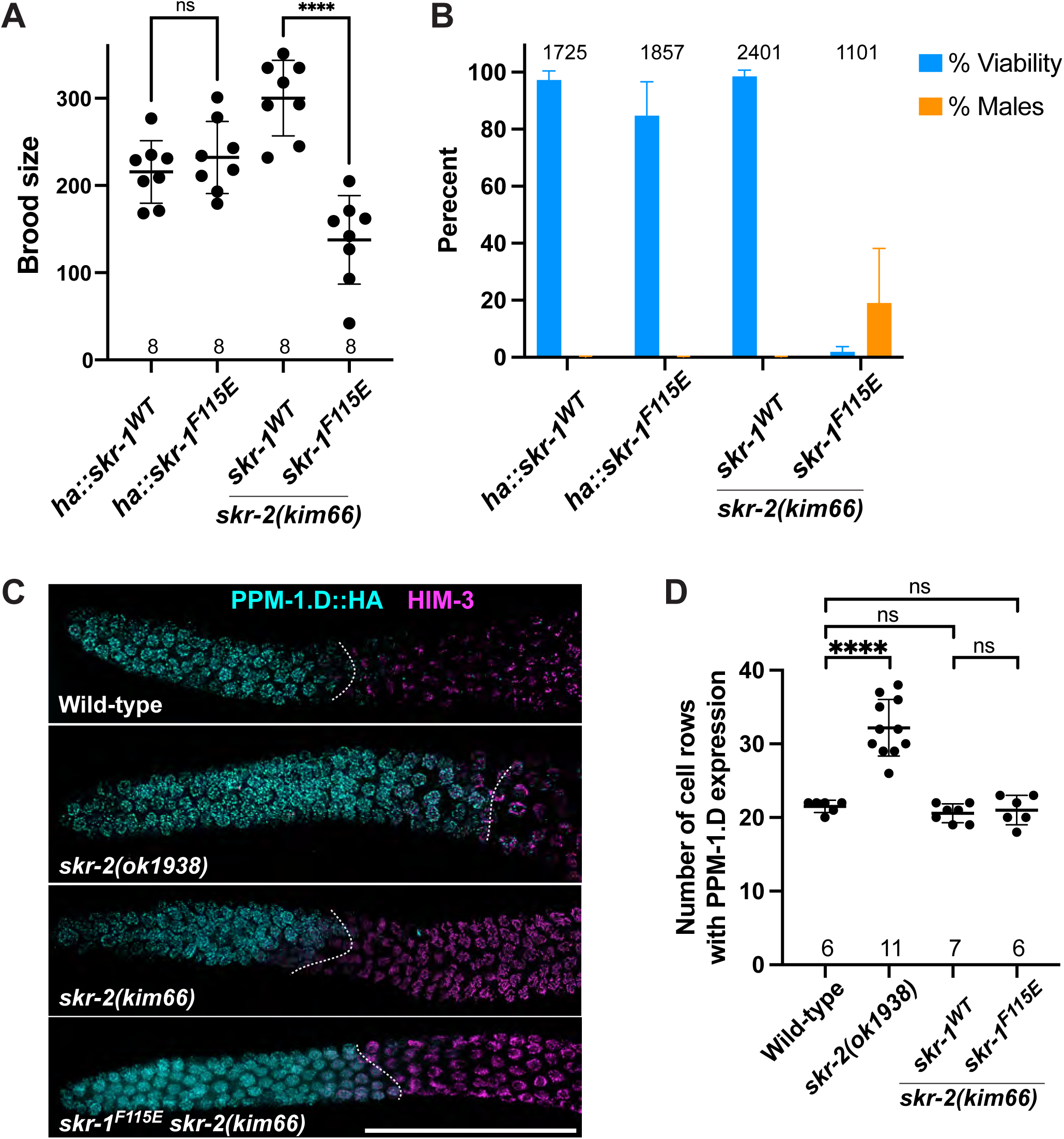
Characterization of the *skr-1* dimer mutant. (A) Graph showing the brood size of indicated genotypes. The numbers of animals scored are shown below the graph. Mean± S.D. is shown; ns, not significant; ****, p<0.0001 by Ordinary one-way ANOVA. (B) Graph showing the percent viable self-progeny and males from indicated genotypes. Mean± S.D. is shown. The number of eggs scored is shown on the top. (C) Composite immunofluorescence images of distal gonads dissected from worm strains harboring indicated *skr-1/2* alleles showing PPM-1.D::HA (cyan) and HIM-3 (magenta) staining. Scale bar, 50 µm. The onset of PPM-1.D degradation is marked by white dotted lines. (D) Graph showing the number of cell rows from the distal tip that displays PPM-1.D expression. Mean ± S.D. is shown, and the numbers of gonads scored are shown below. **** p<0.0001 by Ordinary one-way ANOVA.

**Figure S6.**
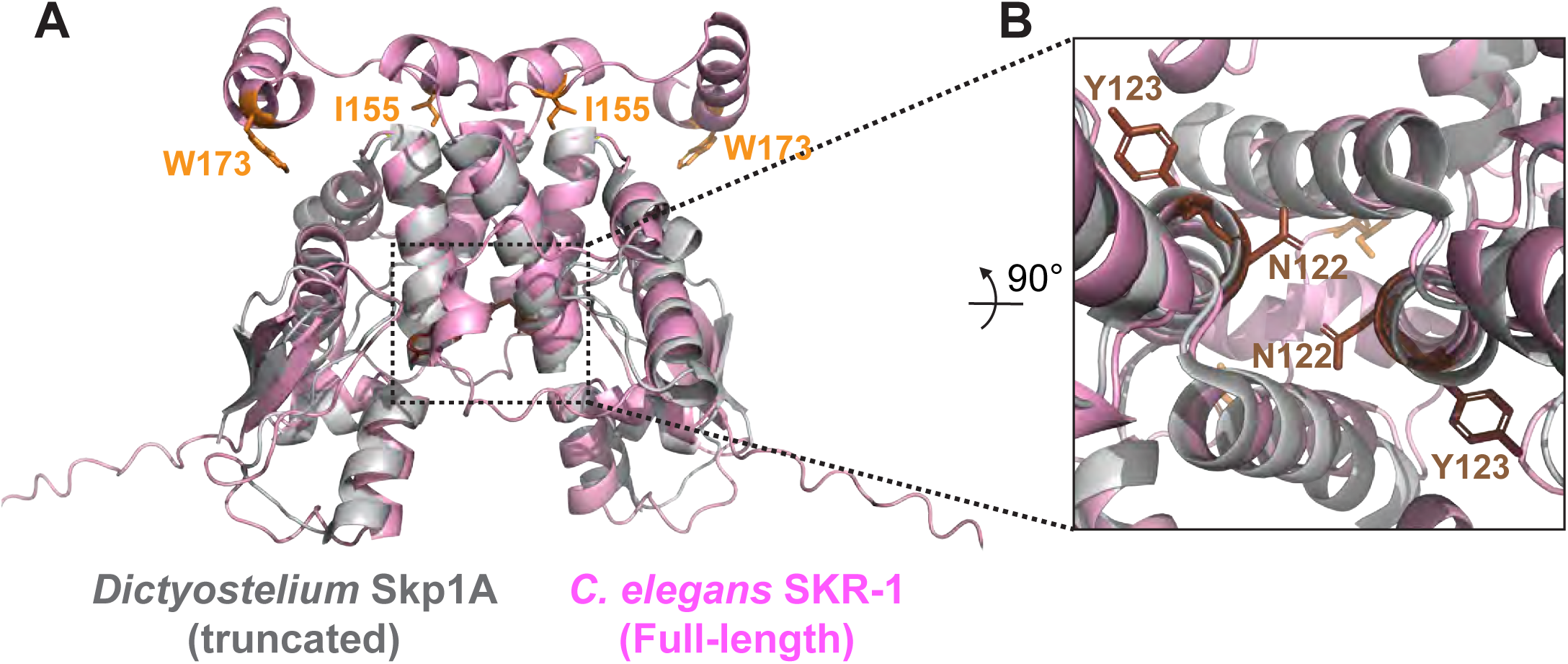
(related to **Figure 6**). **AlphaFold model of SKR-1 dimer highlighting the residues mutated within the CUL-1 or F-box protein-binding interfaces** (A) An AlphaFold model of the full-length *C. elegans* SKR-1 (magenta) superimposed onto the NMR structure of a truncated *Dictyostelium discoideum* Skp1A dimer (grey; PDB, 6V88) with mutated residues are highlighted. (B) The inset shows two residues mutated within the CUL-1-binding interface of SKR-1 (N122 and Y123), which reside within the dimer interface.

**Table S1.**
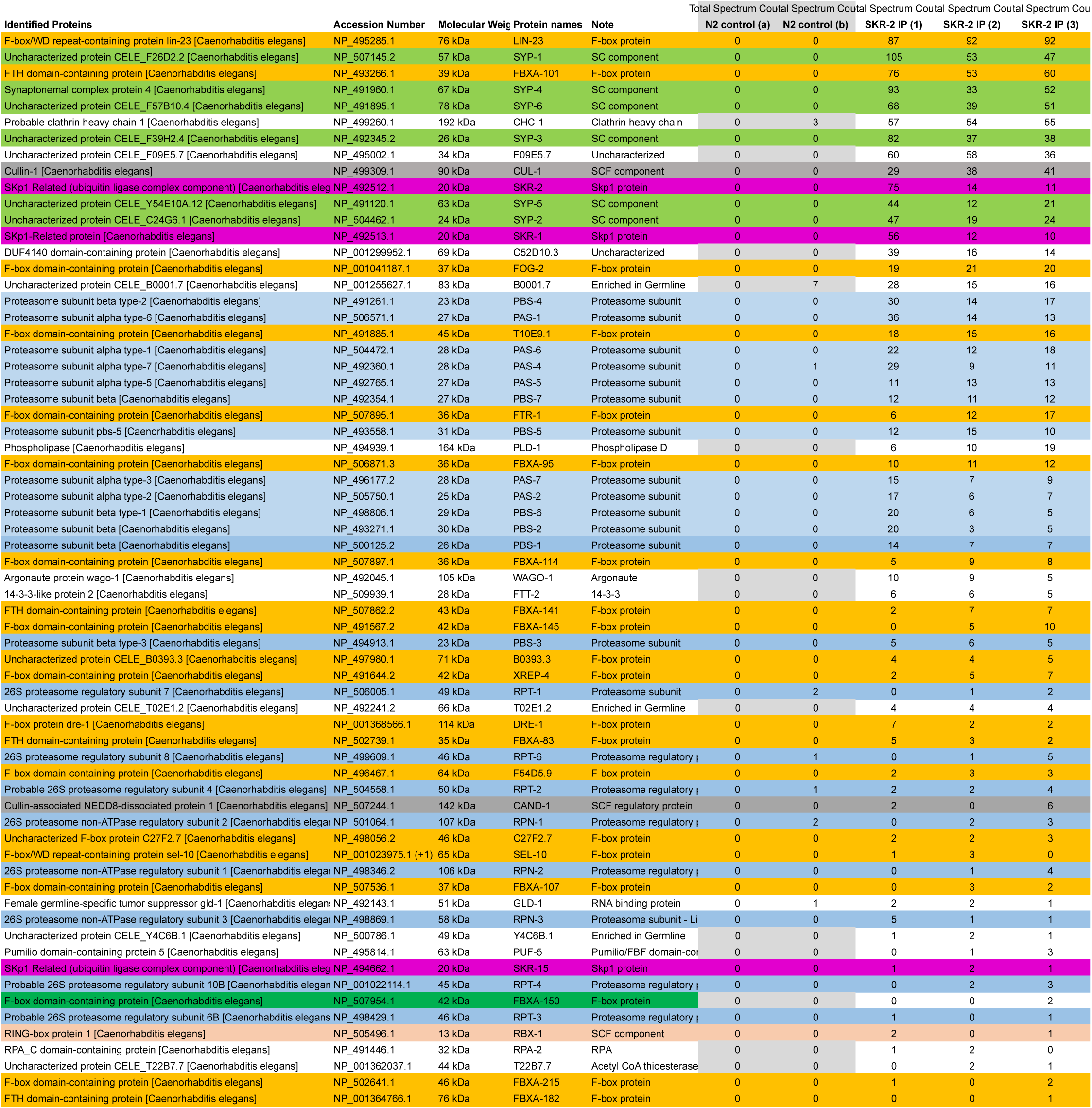
Mass spectrometry analysis of SKR-2 interacting proteins

**Table S2.**
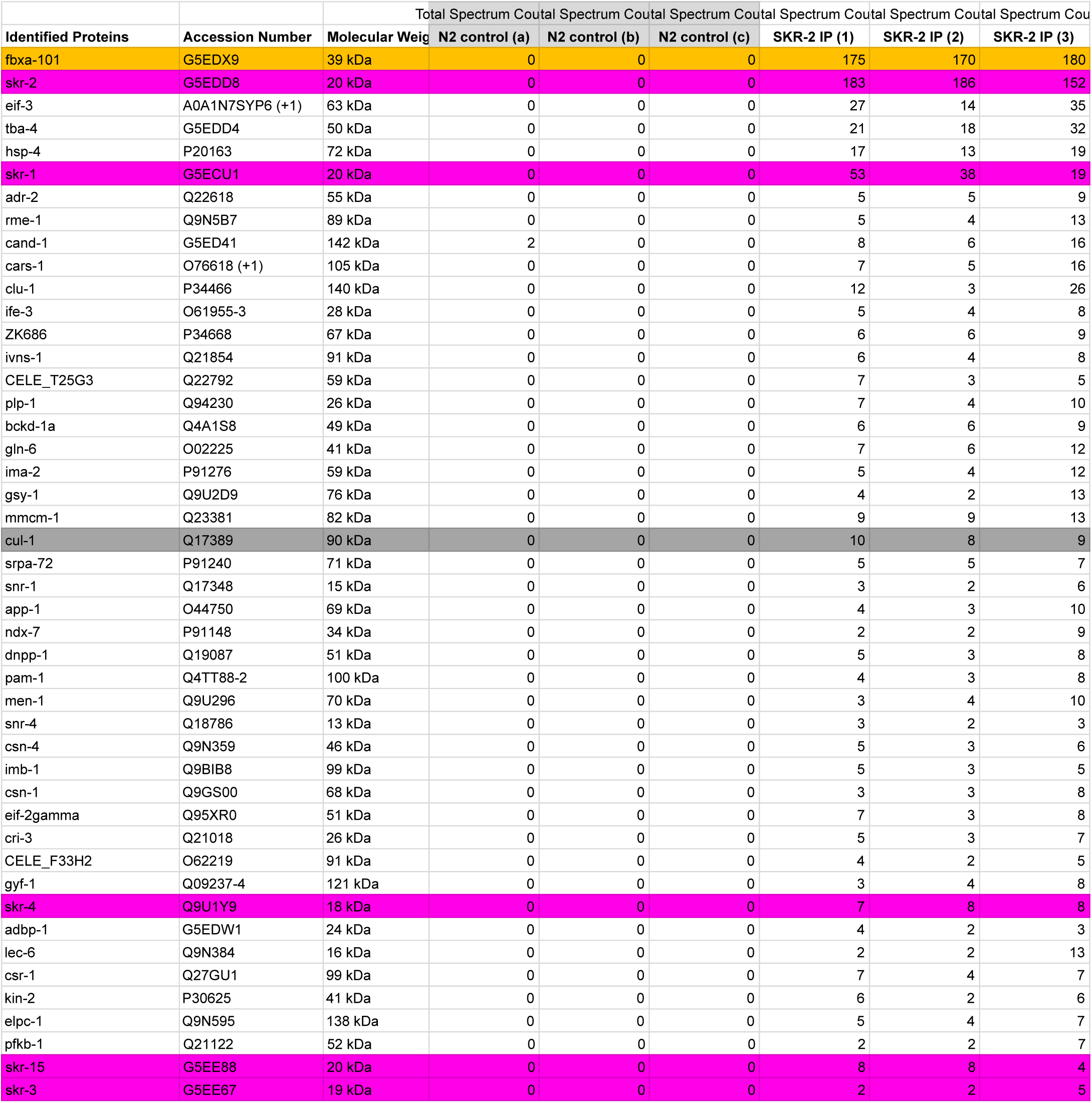
Mass spectrometry analysis of SKR-2^I153A^ ^W171A^ interacting proteins

**Table S3.** Alleles generated in this study

Alleles generated in this study
| Allele | Genotype | Information about mutagenesis |
| --- | --- | --- |
| *kim65* | *skr-2(kim65[skr-2::3flag]) I* | Generated using CRISPR in AV630 |
| *kim66* | *skr-2(kim66[skr-2 Δ50bp]) I* | Generated using CRISPR in AV630 and YKM616 to delete 50 bp in the 1st exon. This results into a premature stop after aa 5. |
| *kim67* | *skr-1(kim67[ha::skr-1]) I* | Generated using CRISPR in N2 |
| *kim68* | *cul-1(kim68[cul-1::ollas]) III* | Generated using CRISPR in AV630 |
| *kim74* | *skr-1(kim74[ha::skr-1(F115E)]) I* | Generated using CRISPR in YKM616 (ha::skr-1) |
| *kim75* | *syp-2(kim75[syp-2::TEV::gfp]) V* | Generated using CRISPR in N2 by InVivo Biosystems |
| *kim76* | *skr-2::3xflag(N120K, Y121K)* | Generated using CRISPR in YKM219 (skr-2::3xflag) |
| *kim77* | *skr-2::3xflag(W171A)* | Generated using CRISPR in YKM219 (skr-2::3xflag) |
| *kim78* | *skr-2::3xflag(I153A, W171A)* | Generated using CRISPR in YKM921 (skr-2::3xflag (W171A)) |

**Table S4.**
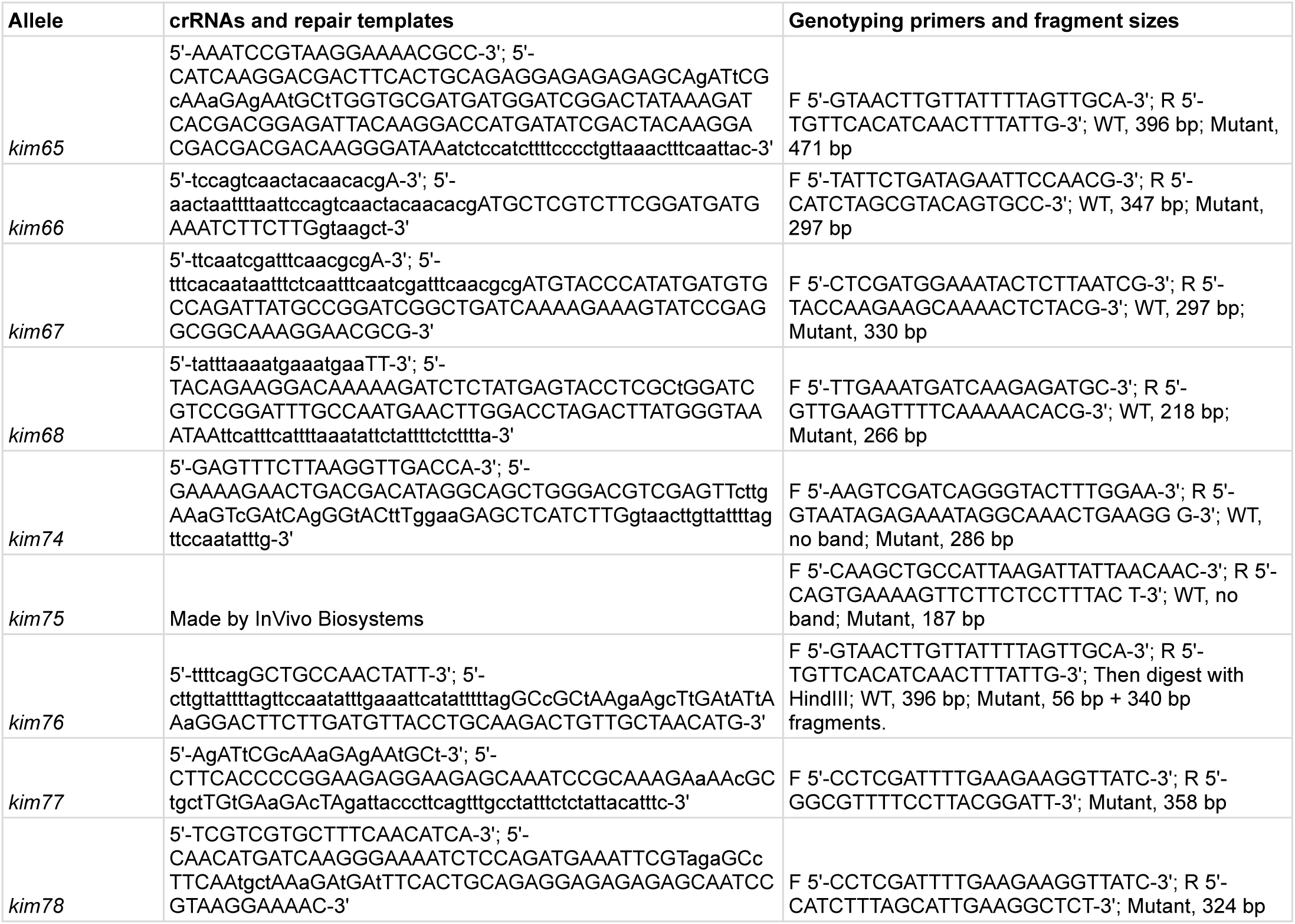
crRNAs, repair templates, and genotyping primers

**Table S5.** Strains used in this study

| Strains | Source | Strain # |
| --- | --- | --- |
| N2 | CGC | CB |
| *mels8[pie-1p::GFP::cosa-1, unc-119(+)] II* | Yokoo et al., 2012 | AV630 |
| *syp-2(ok307) V/nT1[unc-?(n754) let-? qIs50] (IV;V)* | CGC | AV276 |
| *him-3(gk149) IV/nT1[qIs51] (IV;V)* | CGC | AV630 |
| *htp-3(tm3655) I/hT2 [bli-4(e937) let-?(q782) qIs48] (I,III)* | CGC | TY5038 |
| *skr-1(kim67[ha::skr-1]) I* | This study | YKM616 |
| *skr-2(kim65[skr-2::3xflag]) I; mels8[pie-1p::GFP::cosa-1, unc-119(+)] II* | This study | YKM219 |
| *skr-2(kim76[skr-2::3xflag N120K, Y121K]) I; mels8[pie-1p::GFP::cosa-1, unc-119(+)] II* | This study | YKM919 |
| *skr-2(kim65[skr-2::3xflag]) I; syp-2(ok307) V/nT1[unc-?(n754) let-? qIs50] (IV;V)* | This study | YKM643 |
| *skr-2(kim65[skr-2::3xflag]) I; him-3(gk149) IV/nT1[qIs51] (IV;V)* | This study | YKM646 |
| *skr-1(ok1696) I; mels8[pie-1p::GFP::cosa-1, unc-119(+)] II* | This study | YKM217 |
| *skr-1(ok1696) I* | CGC | VC1241 |
| *skr-2(ok1938) I/hT2 [bli-4(e937) let-?(q782) qIs48] (I,III).* | CGC | VC1439 |
| *skr-2(ok1938) I/hT2 [bli-4(e937) let-?(q782) qIs48] (I,III); mels8[pie-1p::GFP::cosa-1, unc-119(+)] II* | This study | YKM627 |
| *skr-2(kim66[skr-2Δ50bp]) I ; mels8[pie-1p::GFP::cosa-1, unc-119(+)] II* | This study | YKM895 |
| *mels8[pie-1p::GFP::cosa-1, unc-119(+)] II; cul-1(kim68[cul-1::ollas]) III* | This study | YKM218 |
| *cul-1(kim68[cul-1::ollas]) III; him-3(gk149) IV/nT1[qIs51] (IV;V )* | This study | YKM868 |
| *chk-2(kim1[3xflag::chk-2]) V; plk-2(kim62[plk-2::gfp]) I; ppm-1.D(kim2[ppm-1.D::HA]) III* | This study | YKM183 |
| *syp-2(kim75[syp-2::TEV::gfp]) V* | This study | YKM853 |
| *htp-3(tm3655) I/hT2 [bli-4(e937) let-?(q782) qIs48] (I,III); syp-2(kim75[syp-2::TEV::gfp]) V* | This study | YKM874 |
| *skr-1(kim74[ha::skr-1(F115E)]) I* | This study | YKM882 |
| *skr-1(kim74[ha::skr-1(F115E)]) skr-2(kim66[skr-2Δ50bp]) I/hT2 [bli-4(e937) let-?(q782) qIs48] (I,III)* | This study | YKM889 |
| *skr-1(kim74[ha::skr-1(F115E)]) skr-2(kim66[skr-2Δ50bp]) I/hT2 [bli-4(e937) let-?(q782) qIs48] (I,III); ppm-1.D(kim2[ppm-1.D::HA]) III* | This study | YKM940 |
| *skr-2(kim76[skr-2::3xflag(N120K, Y121K)]) I; mels8[pie-1p::GFP::cosa-1, unc-119(+)] II* | This study | YKM920 |
| *skr-2(kim77[skr-2::3xflag(W171A)]) I; mels8[pie-1p::GFP::cosa-1, unc-119(+)] II* | This study | YKM921 |
| *skr-2(kim78[skr-2::3xflag(I153A, W171A)]) I; mels8[pie-1p::GFP::cosa-1, unc-119(+)] II* | This study | YKM942 |

| Plasmid name | Backbone | Insert | Note |
| --- | --- | --- | --- |
| pYK407 | pMAL | 6His-SKR-1 (WT) | MBP was deleted |
| pYK412 | pMAL | 6His-SKR-1 (F115E) | MBP was deleted |
| pYK427 | pMAL | 6His-SKR-1 (N122K, Y123K) | MBP was deleted |
| pYK428 | pMAL | 6His-SKR-1 (I155A, W173A) | MBP was deleted |
| pYK423 | pET23c | SYP-2/SYP-1/6His-SYP-3/SYP-4/SYP-5/SKR-1 |  |

## Acknowledgements

We thank Bob Cole in the Mass Spectrometry and Proteomics Facility at Johns Hopkins School of Medicine for mass spectrometry analysis and John Kim (Johns Hopkins University, Baltimore, MD) for the *C. elegans* RNAi libraries. Some strains were provided by the *Caenorhabditis* Genetics Center, which is funded by the National Institutes of Health Office of Research Infrastructure Programs (P40 OD010440). J.H. was supported by the NIH Predoctoral Fellowship (F31HD100142). This work is supported by funding from the European Molecular Biology Laboratory to I. Čavka and S. Köhler and the National Institutes of Health to Y. K. (R35GM124895).

## Author contributions

J.B., B.C., and Y.K. conceived and designed the study. J.B., J.H., and Y.K. generated *C. elegans* strains, performed immunofluorescence, and analyzed the data. J.B. conducted immunoprecipitation of SKR-2 and J.W.B analyzed the synteny of *skr-1/2* genes. J.B., J.W.B., and Y.K. performed RNAi experiments. I.C. and S.K. performed SMLM. B.C. purified recombinant proteins and performed SEC-MALS analyses. Y.K. wrote the paper with input from the other authors.

